# GUN1 influences the accumulation of NEP-dependent transcripts and chloroplast protein import in Arabidopsis cotyledons upon perturbation of chloroplast protein homeostasis

**DOI:** 10.1101/724518

**Authors:** Luca Tadini, Carlotta Peracchio, Andrea Trotta, Monica Colombo, Ilaria Mancini, Nicolaj Jeran, Alex Costa, Franco Faoro, Milena Marsoni, Candida Vannini, Eva-Mari Aro, Paolo Pesaresi

## Abstract

Correct chloroplast development and function require coordinated expression of chloroplast and nuclear genes. This is achieved through chloroplast signals that modulate nuclear gene expression in accordance with the chloroplast’s needs. Genetic evidence indicates that GUN1, a chloroplast-localized pentatricopeptide-repeat (PPR) protein with a C-terminal Small MutS-Related (SMR) domain, is involved in integrating multiple developmental and stress-related signals in both young seedlings and adult leaves. Recently, GUN1 was found to interact physically with factors involved in chloroplast protein homeostasis, and with enzymes of tetrapyrrole biosynthesis in adult leaves that function in various retrograde signaling pathways. Here we show that, following perturbation of chloroplast protein homeostasis i) by growth in lincomycin-containing medium, or ii) in mutants defective in either the FtsH protease complex (*ftsh*), plastid ribosome activity (*prps21-1* and *prpl11-1*) or plastid protein import and folding (*cphsp70-1*), GUN1 influences NEP-dependent transcript accumulation during cotyledon greening and also intervenes in chloroplast protein import.

## INTRODUCTION

Chloroplast biogenesis is achieved through a cascade of events that include cytosolic synthesis of chloroplast-targeted proteins, followed by their import, assembly and folding, while unfolded/misfolded proteins and damaged or malformed chloroplasts are degraded (Sakamoto et al., 2008; Jarvi and López-Jauez, 2013; Izumi and Nakamura, 2018; Otegui, 2018). Most chloroplast-targeted proteins enter the chloroplasts via the protein complexes TOC (‘translocon at outer envelope of chloroplast’) and TIC (‘translocon at inner envelope of chloroplast’). On the cytosolic side, precursor proteins are directed to the Toc34 and Toc159 receptors through the interaction of their chloroplast-targeting peptides with the chaperone heat shock protein 90 (Hsp90) and the guidance complex formed by Hsp70 and a 14-3-3 protein. Active transport through the chloroplast envelope is then mediated by an ATP-dependent import motor, consisting of Hsp70, Hsp90 and the stromal Hsp93 (ClpC2) proteins (Kessler and Schnell, 2009; Sjuts et al., 2017).

On the stromal side, transcription of the plastid-encoded genes in higher plants requires two different RNA polymerases: the nuclear-encoded RNA polymerase (NEP), also termed RpoTp (located specifically in the chloroplasts and encoded by the *RPOT3* gene) or RpoTmp (located both in mitochondria and plastids and encoded by the *RPOT2* gene) and the plastid-encoded RNA polymerase (PEP) (Yu et al., 2014; Börner et al., 2015). NEP is a monomeric T3–T7 bacteriophage-type RNA polymerase that is mainly responsible for transcribing housekeeping genes, whereas PEP is a bacterial-type multisubunit enzyme largely tasked with transcribing photosynthesis-related genes. Chloroplast development is associated with a shift in the primary RNA polymerase from NEP to PEP. Furthermore, compensatory responses can be observed between the two RNA polymerases, as depletion of PEP activity increases the expression of NEP-dependent transcripts (Myouga et al., 2008; Pfalz et al., 2006; Ramos-Vega et al., 2015; Zhou et al., 2009). This process was designated as the “Δ-*rpo* phenotype” (Allison et al., 1996; Hajdukiewicz et al., 1997; De Santis-Maciossek et al., 1999), but the underlying mechanisms are still unknown.

Both imported proteins and proteins synthesized in the chloroplast need to be folded and further processed, or degraded if misfolded. Chloroplast protein folding is mediated by chaperones, including Hsp70, Cpn60 and ClpB3, while protein degradation is accomplished by the stromal protease Clp, a multimeric complex made up of chaperones and serine protease subunits, which serves general housekeeping functions (Flores-Pérez and Järvis, 2013; Nishimura et al., 2016; Nishimura et al., 2017). In addition, the thylakoid-associated heteromeric FtsH proteases, comprised of FtsH1 or FtsH5 (type A) and FtsH2 or FtsH8 (type B) subunits, participate in the Photosystem II (PSII) repair cycle, together with the DEG serine proteases (Kato and Sakamoto, 2018).

Besides transcription, translation and post-translational processes, chloroplast biogenesis also requires tight coordination of the transcription of thousands of nuclear genes with the expression of the relatively few plastid genes in order to meet the needs of the developing chloroplasts. This coordination is achieved through extensive exchange of information between the plastids and the nucleus via, for instance, biogenic retrograde signaling (Pogson et al., 2008; Woodson and Chory, 2008; Chan et al., 2016). In particular, *GUN1*, which codes for a chloroplast pentatricopeptide repeat (PPR) protein, has been shown to have a role in plastid gene expression (PGE; Tadini et al., 2016) and to act as an integrator of multiple retrograde signals. The exact molecular role of GUN1 has long remained enigmatic, but the recent identification of a set of GUN1-interacting proteins in leaf tissue of Arabidopsis, together with the observation that the DEAD-box RNA helicase RH50, associated with plastid ribosome functioning, is functionally related to GUN1, point to a role for GUN1 in maintenance of protein homeostasis (reviewed in Colombo et al., 2016; Tadini et al., 2016; Paieri et al., 2018). Nevertheless, a detailed molecular analysis that could uncover the precise mode of action of the GUN1 protein in Arabidopsis cotyledons is still lacking. Here, we have investigated the role of GUN1 under conditions that trigger its accumulation to detectable levels, *i.e.* in Arabidopsis cotyledons during early stages of chloroplast biogenesis in the presence of either lincomycin (Lin; Wu et al., 2018) or of mutations that affect plastid protein homeostasis, by depleting levels of the FtsH protease complex (*ftsh*), reducing plastid ribosome activity (*prps21-1*, *prpl11-1* and *rh50-1*) or disrupting plastid protein import and folding (*cphsp70-1*). We show that GUN1 is important for NEP-dependent transcript accumulation, possibly being a key component of the NEP-dependent compensatory mechanism activated upon conditions that alter chloroplast protein homeostasis and PEP activity.

## RESULTS

### GUN1 influences the accumulation of NEP-dependent transcripts in response to perturbation of chloroplast protein homeostasis

As a member of the PPR protein family, a possible role for GUN1 as either a key regulator of plastid transcript synthesis or post-transcriptional processing can be envisaged to explain the additive phenotypic effects observed in *gun1-102 prpl11-1*, *gun1-102 prps17-1* and *gun1-102 prpl24-1* double mutant seedlings reported in Tadini et al. (2016) and Paieri et al. (2018) and the similar patterns of epistatic effects evoked by *rh50-1* and *gun1-102* mutations (Paieri et al., 2018). To substantiate this idea, we used RNA gel-blot hybridization to measure (at 6 days after sowing, DAS) the levels of NEP- and PEP-dependent plastid transcripts in cotyledons of Col-0, *gun1-102* and other mutants affected at different levels in chloroplast protein homeostasis, and grown in the absence or presence of lincomycin (Figure 1; Figure S1 and Table S1). Transcripts encoding the large subunit of RUBISCO (*rbcL*), the *α* subunit of ATPase (*atpα*), the A subunit of PSI core (*psaA*) and the D1 subunit of PSII reaction center (*psbA*) were selected as typical products of PEP. Instead, probes specific for *Tic214* (*Ycf1*, which encodes a constituent of the 1-MDa TIC complex), the 3’ portion (*rps12-3’*) of *rps12* (which codes for the S12 subunit of plastid ribosomes), the *rpoA* and *rpoB* sequences that code for the α and β subunits of the PEP RNA polymerase, respectively, and *clpP1* which encodes the proteolytic subunit 1 of the ATP-dependent Clp protease were selected to monitor the accumulation of NEP-dependent transcripts. Total RNA extracted from Col-0 and *gun1-102* cotyledons grown in the absence or increasing concentrations of lincomycin were employed in the first hybridization assay (Figure 1a and Table S1). In particular, the expected increase in NEP-dependent transcripts (*rpoA* and *rps12*) in Col-0 cotyledons was already detectable at minimal concentrations of lincomycin (5.5 μM) in the growth medium, when the inhibitory effect on cotyledon greening was barely discernible (Figure 2). In the case of *rpoA* mRNAs, the increase in the total transcript amount was proportional to the increase in the lincomycin concentration and the progressive impairment of cotyledon greening, which reaches its maximum at 550 μM lincomycin, when plastid rRNAs become undetectable with methylene blue (see Figure 1a) and the proplastid-to-chloroplast transition is completely suppressed (Figure 2). A similar transcript accumulation pattern in response to different lincomycin concentrations was observed in *gun1-102* cotyledons, although the total levels of *rpoA* and *rps12-3’* transcripts were markedly reduced, possibly explaining the higher sensitivity of *gun1-102* seedlings to minimal concentration of lincomycin (5.5 μM), compared to Col-0 (Figure 2). To obtain independent evidence for the specific role of GUN1 in NEP-dependent transcription accumulation, total RNA was also isolated from cotyledons of Arabidopsis mutants lacking the S21 (*prps21-1*) and L11 subunits (*prpl11-1*) of plastid ribosomes (Pesaresi et al., 2001; Morita-Yamamuro et al., 2004), and from the corresponding *gun1-102 prps21-1* and *gun1-102 prpl11-1* double mutants (Tadini et al., 2016; Figure 1b). RNA gel-blot hybridizations confirmed the substantial increase in levels of mRNAs synthesized by the NEP in *prps21-1* and *prpl11-1*, but not in *gun1-102 prps21-1* and *gun1-102 prpl11-1* cotyledons (Table S1). The specific involvement of GUN1 protein in NEP-dependent transcript accumulation was further verified by using Arabidopsis mutants altered in chloroplast protein homeostasis such as *ftsh5-*3, lacking the FtsH protease complex, *cphsp70-1*, affected in plastid protein import and folding (Flores-Pérez and Jarvis, 2013; Sjuts et al., 2017), or *rh50-1*, reported to be functionally related to GUN1 and characterized by altered plastid rRNA maturation (Paieri et al., 2018). No changes in the selected NEP-dependent mRNAs were observed in *gun1-102* and *gun1-102 ftsh5-3* cotyledons (relative to Col-0), while all of them exhibited a slight increase in level in *ftsh5-3* seedlings (Figure 1c). In contrast, levels of NEP-dependent transcripts increased substantially in Col-0 and *ftsh5-3* cotyledons grown on MS medium supplemented with lincomycin, but not in *gun1-102* or *gun1-102 ftsh5-3* cotyledons (Figure 1c). Similarly, the abundance of NEP-dependent transcripts such as *rpoA* and *rps12-3’* was also substantially increased in *cphsc70-1* cotyledons and reduced to Col-0 levels in *gun1-102 cphsc70-1* seedlings (Figure 1d). Finally, the *rh50-1* mutant behaved in a way similar to Col-0 in the presence of lincomycin with respect to NEP-dependent transcript accumulation, but almost no compensatory response could be observed in *gun1-102* (Figure 1e), implying that the increase in NEP-related transcripts upon perturbation of plastid protein homeostasis is a GUN1-specific process.

**Figure 1.**
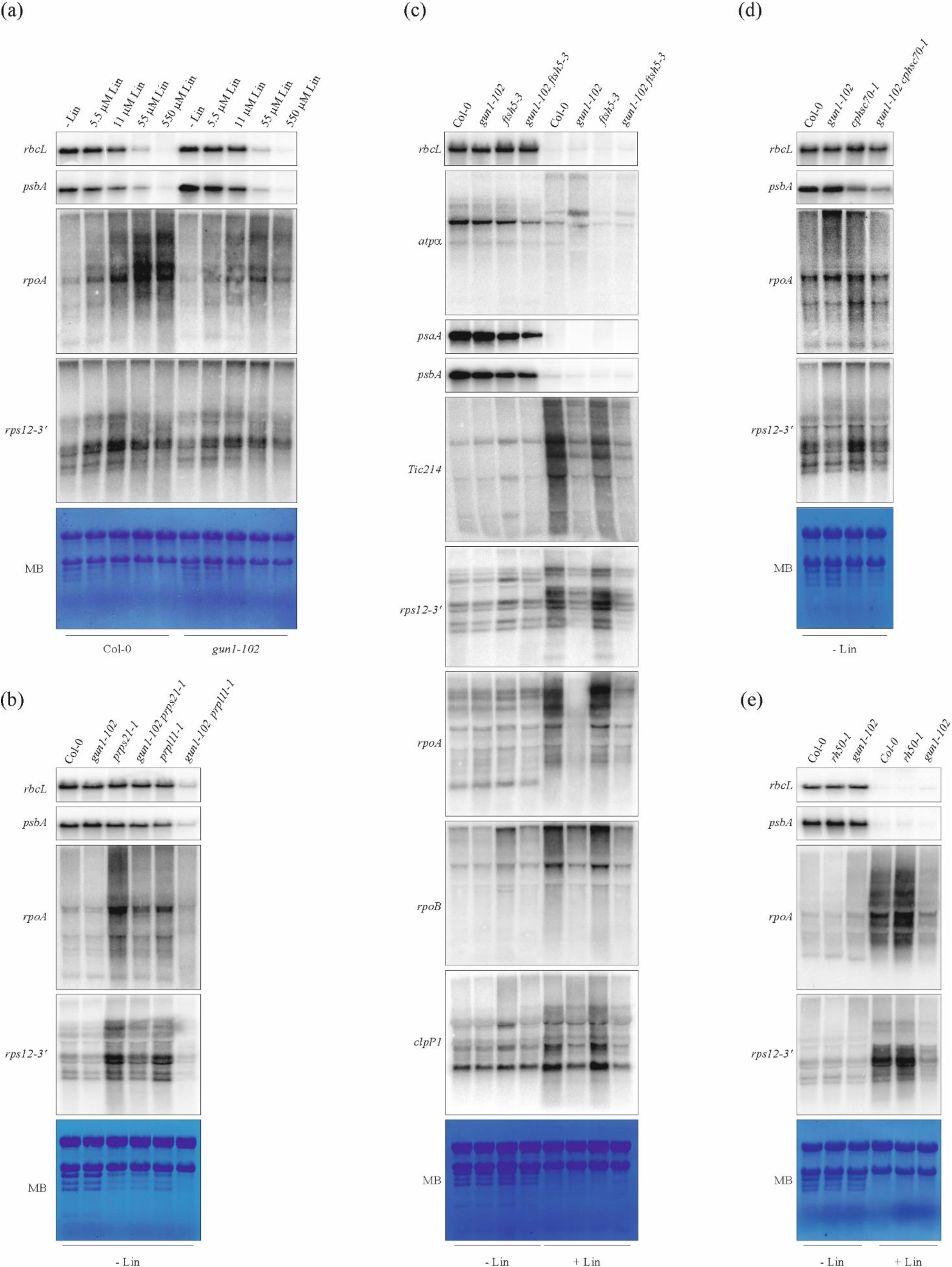
Expression of selected transcripts in Col-0 and in mutant seedlings altered in chloroplast protein homeostasis. **(a)** Gel-blot analysis of total RNA isolated from cotyledons of Col-0 and *gun1-102* seedlings grown in either the absence (-Lin) or presence of different concentrations of lincomycin (5.5 μM, 11 μM, 55 μM, 550 μM; see also Figure 2). **(b)** Gel-blot analysis of total RNA isolated from cotyledons of Col-0, *gun1-102*, *prps21-1* and *prpl11-1* (mutants defective in protein synthesis in plastids), together with *gun1-102 prps21-1* and *gun1-102 prpl11-1* double mutants. **(c)** Gel blot analysis of total RNA isolated from cotyledons of Col-0, *gun1-102*, *ftsh5-3* (mutant defective in protein degradation) and *gun1-102 ftsh5-3* grown on MS medium with and without Lin. **(d)** Gel-blot analysis of total RNA isolated from cotyledons of Col-0, *gun1-102*, *cphsp70-1* (mutant defective in protein import and protein folding) and *gun1-102 cphsp70-1* seedlings. (**e**) Gel blot analysis of total RNA isolated from cotyledons of Col-0, *rh50-1* (mutant defective in in rRNA maturation) and *gun1-102* grown om MS medium with and without Lin. Probes specific for the PEP-dependent *rbcL*, *psaA*, *psbA*, *atpα* and NEP-dependent *Tic214* (*Ycf1*), *rpoA*, *rpoB*, *clpP1* and *rps12-3’* plastid RNAs were used for hybridization. A representative result from three independent experiments is shown. The methylene blue (MB)-stained filters are shown as the loading control.

**Figure 2.**
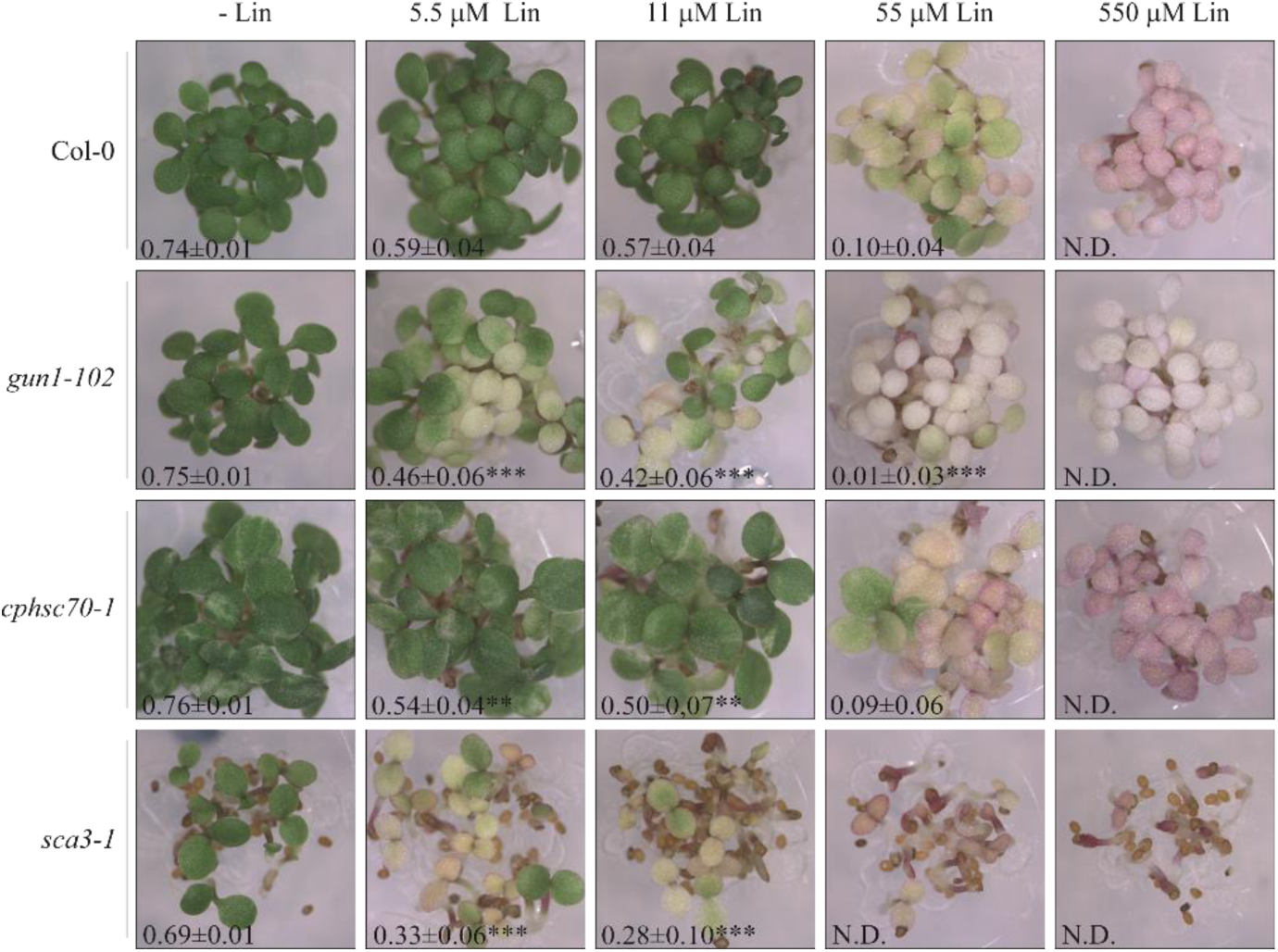
Visible phenotypes and photosynthetic performance of Col-0, *gun1-102, sca3-1* and *cphsc70-1* seedlings at 6 DAS, grown on MS medium in the absence (-Lin) or presence of different concentrations of lincomycin (5.5 μM, 11 μM, 55 μM, 550 μM). The effective quantum yield of PSII (Φ_II_) was measured by using the IMAGING-PAM (WALZ) and reported as average ± standard deviation of five independent measurements. Clearly, seedlings devoid of GUN1 (*gun-102*) and RpoTp (*sca3-1*) proteins show enhanced sensitivity to lincomycin. Asterisks indicate the statistical significance, with respect to Col-0, of Φ_II_ values, as evaluated by Student’s t-test and Welch correction (*, p<0,05; **, p<0,01; ***, p<0,001).

### GUN1 physically interacts with RpoTp and favors its activity upon depletion of PEP

The plastid gene expression profile reported in Figure 1 (see also Table S1) suggests that GUN1 is required for the expression of genes transcribed by NEP upon depletion of PEP activity. To investigate if the increased steady-state levels of the NEP-dependent transcripts observed in Col-0 compared to *gun1-102* plastids in presence of lincomycin was due to increased transcriptional activity rather than posttranscriptional effects such as decreased RNA stability, run-on transcription assays for the NEP-dependent transcripts *rpoA* and *rps12*, and the PEP-dependent *rbcL*, *psaA* and *psbA* RNAs were performed in plastids isolated from Col-0, *gun1-102* and plastids of *gun1-102* in the presence of either the purified GUN1 recombinant protein (+ recGUN1) or RH50 recombinant protein (+ recRH50 - Dead-box RNA Helicase RH50; see also Paieri et al., 2018), the latter used as control (Figure 3a,b and Figure S2a,b). This assay demonstrated that the levels of newly synthesized NEP-dependent transcripts were significantly higher in Col-0 and *gun1-102* + recGUN1 plastids compared to *gun1-102*, whereas the PEP-dependent transcripts accumulated to similar levels (Figure 3a,b and Figure S2a,b), indicating that GUN1 is required for RopTp transcriptional activity rather than influencing RNA stability. This is also supported by the fact that no major differences could be observed in the accumulation of the RpoTp enzyme in both Col-0 and *gun-102* seedlings (+/- Lin), as revealed by immunoblot assay (Figure 3c).

**Figure 3.**
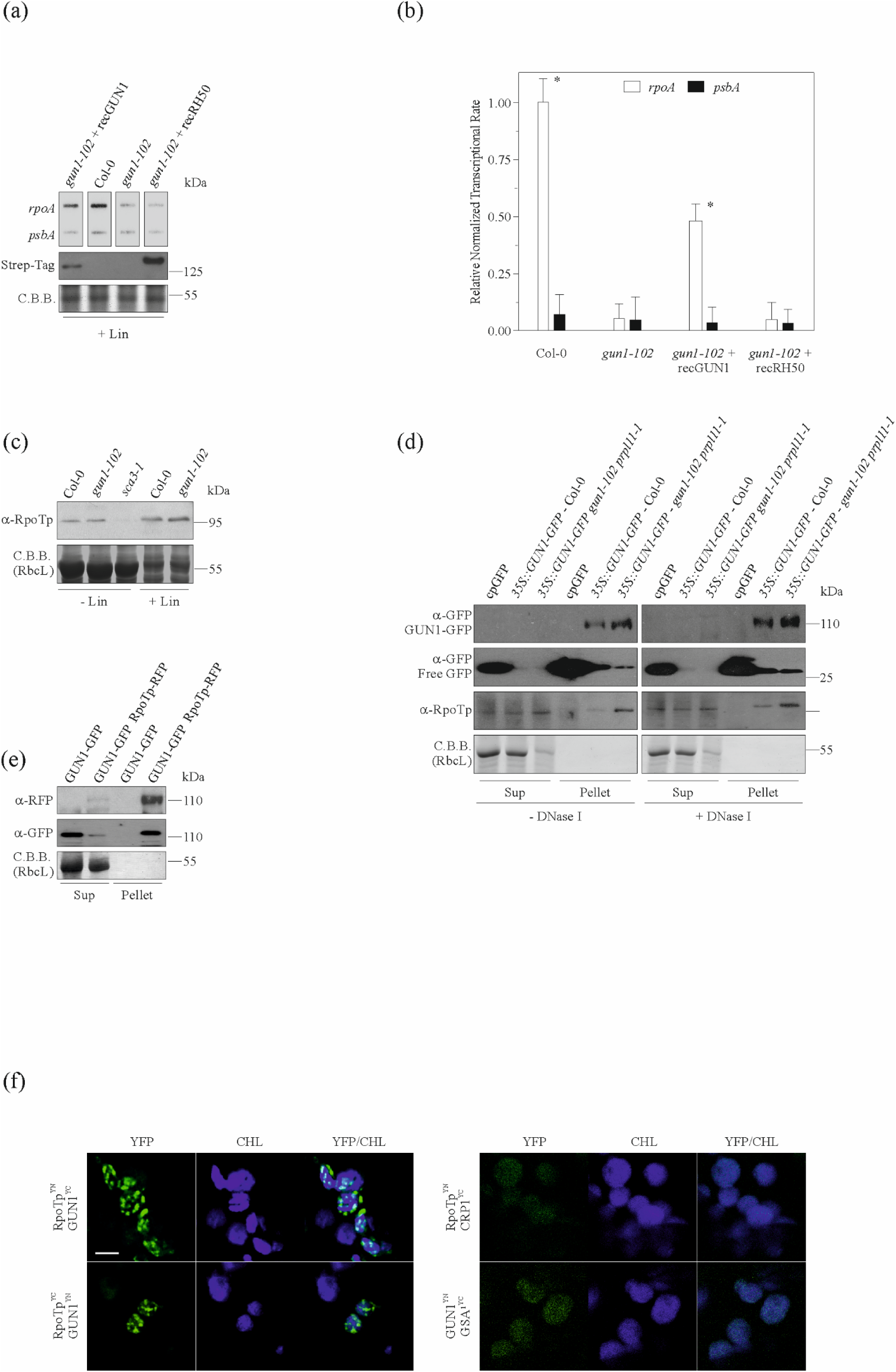
The role of GUN1 in NEP-dependent transcriptional activity. **(a)** Run-on transcription assay in Col-0, *gun1-102*, *gun1-102* + recGUN1 and *gun1-102* + recRH50 plastids isolated from seedlings grown on MS medium with Lin. Filters with *rpoA* (NEP-dependent gene) and *psbA* (PEP-dependent gene) probes were hybridized with total [^32^P]-RNA from Col-0, *gun1-102*, *gun1-102* + recGUN1 and *gun1-102* + recRH50 isolated plastids. A representative result from three independent experiments is shown. An immunoblot with the Strep-tag specific antibody is shown to verify the presence of GUN1 and RH50 recombinant proteins in the reaction mixture. The C.B.B stained nylon membrane, corresponding to RbcL migration region, was used to verify equal loading. (**b**) Quantification of Run-on signals in (a). Data are the mean ± s.d. from quantification of three independent experiments. Asterisks indicate the statistical significance, as evaluated by Student’s t-test (p<0.05). (**c**) Immunoblot analysis of total proteins isolated from Col-0 and *gun1-102* cotyledons grown on MS medium with and without lincomycin (-/+ Lin) to monitor RpoTp accumulation. The *sca3-1* mutant, carrying a single nucleotide exchange in the At2g24120 gene encoding the RpoTp enzyme (Hricová et al., 2006; see also Figure S1), has been used to check for the RpoTp antibody specificity. (**d**) Immunoblot analysis of protein fractions obtained from immunoprecipitation experiments using the anti-GFP antibody carried out in cpGFP, (Col-0 seedlings carrying a chloroplast-located GFP), *35S::GUN1-GFP* in Col-0 and *35S::GUN1-GFP* in *gun1-102 prpl11-1* Arabidopsis cotyledons. Equal volumes of supernatant (Sup) and Pellet preparations in either the absence or presence of DNase I (+/-DNAase I) were loaded onto SDS-PAGEs. The pellets from 35S::GUN1-GFP immunoprecipitations gave very strong signals with respect to the corresponding supernatants, implying quantitative precipitation of GUN1-GFP protein. The anti-RPOTp antibody was able to detect the presence of the nuclear-encoded RNA polymerase in the corresponding pellet fractions, but not in cpGFP pellet, indicating that GUN1 interacts physically with RpoTp. The C.B.B stained nylon membrane, corresponding to RbcL migration region, was used to verify equal loading. Note that free GFP signals in *35S::GUN1-GFP* in Col-0 and *35S::GUN1-GFP* in gun1-102 prpl11-1 pellets are the results of protein degradation and/or products of alternative translation initiation (**e**) Immunoblot analysis of protein fractions obtained from immunoprecipitation experiments using the anti-GFP antibody from GUN1-GFP and RpoTp-RFP protein chimeras, transiently (co)expressed in young tobacco leaves. Equal volumes of supernatant (Sup) and Pellet preparations were loaded onto the SDS-PAGE. The pellet from GUN1-GFP-RpoTp-RFP coimmunoprecipitation gave very strong signals with respect to GFP and RFP specific antibodies, implying the existence of a GUN1-RpoTp protein complex. The GUN1-GFP construct alone was used as negative control of the immunoprecipitation assay. The C.B.B. stained SDS-PAGE, corresponding to the RbcL migration region, was used to verify equal loading. (**f**) BiFC assay in tobacco leaves detected by fluorescence confocal microscopy. GUN1 and RPOTp were fused to the C-terminal (YC) and the N-terminal (YN) ends of the Venus protein, respectively, and cotransformed into tobacco leaves. The reciprocal chimer combination was tested, too. In both cases, representative images of YFP fluorescence reconstitution (signaling positive interaction), chlorophyll autofluorescence (CHL), and their overlay (YFP/CHL) are shown. The chloroplast nucleoid located PPR protein, CRP1, in combination with RpoTp and GUN1^YN^-GSA1^YC^ were used as negative controls. Scale bars = 5 μm.

To explore the existence of physical interactions between GUN1 and the RpoTp enzyme, total protein extracts from Arabidopsis Col-0 and *gun1-102 prpl11-1* cotyledons expressing GUN1-GFP under the control of the *35S-CaMV* promoter (*35S::GUN1-GFP* – Col-0, *35S::GUN1-GFP* – *gun1-102 prpl11-1*) were isolated and the fusion protein was immunoprecipitated using an anti-GFP serum (Figure 3d). As a control, we performed mock precipitations with total protein extracted from Arabidopsis Col-0 cotyledons, carrying a chloroplast-located GFP protein (cpGFP), using the same GFP antibody. Interestingly, the increased abundance of GUN1-GFP in the immunoprecipitated fraction (PELLET) of *35S::GUN1-GFP* – *gun1-102 prpl11-1* sample with respect to *35S::GUN1-GFP* – Col-0, confirmed the increased stability of GUN1 protein under conditions that alter plastid protein synthesis, as reported in Wu et al. (2018; see also Figure S2c). Furthermore, the RpoTp enzyme could be exclusively detected in the immunoprecipitated fractions (Pellet) of *35S::GUN1-GFP* – Col-0 and *35S::GUN1-GFP* – *gun1-102 prpl11-1* samples either in the presence or absence of DNase I (+/- DNase I) by immunoblot, demonstrating that GUN1 and RpoTp proteins are part of the same protein complex and that chloroplast DNA is not required for the establishment of protein interaction. The existence of the GUN1-RpoTp protein complex was further proved by co-immunoprecipitation of GUN1-GFP and RpoTp-RFP protein chimeras, transiently coexpressed in young tobacco leaves (Figure 3e). In addition, BiFC assays in tobacco leaf mesophyll cells corroborated the interactions of GUN1 with RpoTp (Figure 3f). In particular, the distribution of yellow fluorescence signals resulting from this protein-protein interaction was localized to distinct spots within chloroplasts, resembling the distribution of green fluorescence emitted by the GUN1-GFP construct observed in Koussevitzky et al. (2007) and Tadini el al. (2016). The combination of CRP1^YC^, a PPR protein associated to chloroplast nucleoids (Ferrari et al., 2017), with RpoTp^YN^, as well as GUN1^YN^-GSA1^YC^ (Glu-1-semialdehyde 2,1-aminomutase; see also Tadini et al., 2016), used as negative controls, failed to produce the YFP signal, supporting further the specificity of GUN1-RpoTp interaction. In summary, GUN1 interacts with RpoTp possibly increasing the NEP-dependent chloroplast transcription rate upon depletion of PEP activity.

### The absence of GUN1 alters the chloroplast translocon machinery

Loss of GUN1 function, under conditions that affect chloroplast protein homeostasis, is expected to severely perturb chloroplast proteome due to GUN1-dependent modulation of RpoT activity and the major role of GUN1 in the chloroplast-to-nucleus retrograde communication. In particular, immunoblot analyses of total protein extracts from Col-0 and *gun1-102* cotyledons subjected to lincomycin inhibition of plastid protein synthesis showed no accumulation of the antenna proteins of PSI (Lhca1-to-Lhca4) and PSII (Lhcb1, Lhcb2, Lhcb4 and-Lhcb5; Figure 4a and Table 1 for the quantification of protein amounts), and of the small (RbcS, Figure 4a) or large (RbcL) subunits of RUBISCO (see the C.B.B.-stained gel: RbcL, the most abundant band migrating at around 55 kDa in the control samples is mostly absent in lincomycin-treated seedlings), whereas identical protein amounts accumulated in Col-0 and *gun1-102* cotyledons grown on MS medium without lincomycin.

**Figure 4.**
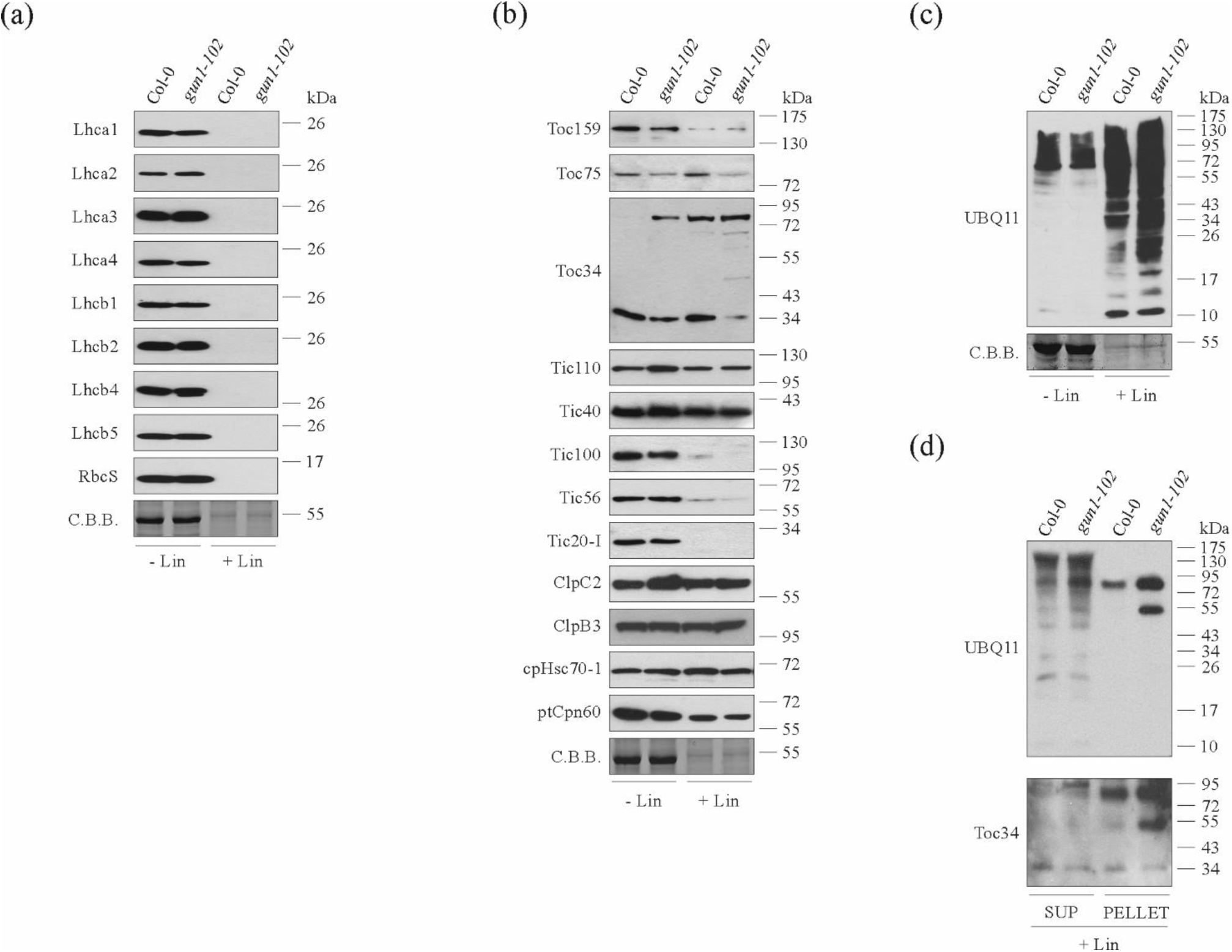
Immunoblot analyses in Col-0, *gun1-102* cotyledons grown on MS medium with and without lincomycin. **(a)** Immunoblot analyses of total protein extracts for the antenna subunits of photosynthetic complexes (Lcha1-Lhca4, Lhcb1-Lhcb2, Lhcb4-Lhcb5), and the small subunit of RUBISCO (RbcS). Plant material corresponding to 5 mg of seedling fresh weight, collected at 6 DAS, was used for the analyses. **(b)** Immunoblot analyses of components of the Tic and Toc translocation machinery, together with the members of the stromal chaperone system. Nitrocellulose filters bearing fractionated cotyledon total proteins were probed with antibodies directed against subunits of the TOC (Toc159, Toc75, and Toc34), TIC (Tic110, Tic40) and 1-MDa TIC (Tic100, Tic56, Tic20-I) complexes. Furthermore, antibodies specific for elements of the stromal chaperone system responsible of protein import and protein folding were also used (ClpC2, ClpB3, cpHsp70-1, ptCpn60). **(c)** Ubiquitination levels in total cotyledon protein extracts from Col-0 and *gun1-102* seedlings grown on MS medium with and without lincomycin, detected with the UBQ11-specific antibody. **(d)** Immunoblot analysis of protein fractions from Col-0 + Lin and *gun1-102* + Lin cotyledons isolated by immunoprecipitation with the anti-Toc34 antibody. Equal volumes of supernatant and pellet preparations were loaded onto the gels, and replicate filters were probed with UBQ11- and Toc34-specific antibodies. The protein bands migrating between 50 and 90 kDa in *gun1-102* and Col-0 samples, recognized by the UBQ11 antibody, were also detected with the Toc34 antibody, demonstrating that these two bands represent ubiquitinated forms of Toc34. The C.B.B. stained SDS-PAGEs, corresponding to RbcL migration region, were used to verify equal loading.

**Table 1.** Quantification of proteins in light-adapted cotyledons of Col-0 and *gun-102* seedlings grown on MS medium with and without lincomycin (+/- Lin). Col-0 levels are set to 1 (100%). Values are means ± SD from three independent protein gel blots (see Figure 4a,b). Asterisks indicate the statistical significance, as evaluated by Student’s t-test and Welch correction (*, p<0.05; **, p<0.01; ***, p<0.001). ^a^Abundance of proteins migrating at higher molecular weights as observed in the case of Toc34 subunit. nd, not detected.

| Protein | *gun1-102* | Col-0 + Lin | *gun1-102* + Lin |
| --- | --- | --- | --- |
| Lhca1 | 0.91 ± 0.11 | nd | nd |
| Lhca2 | 1.08 ± 0.07 | nd | nd |
| Lhca3 | 1.11 ± 0.12 | nd | nd |
| Lhca4 | 0.88 ± 0.12 | nd | nd |
| Lhcb1 | 0.94 ± 0.08 | nd | nd |
| Lhcb2 | 0.89 ± 0.04 | nd | nd |
| Lhcb4 | 1.02 ± 0.09 | nd | nd |
| Lhcb5 | 1.06 ± 0.08 | nd | nd |
| RbcS | 1.05 ± 0.12 | nd | nd |
| Toc159 | 0.59 ± 0.04** | 0.11 ± 0.01 | 0.14 ± 0.01** |
| Toc75 | 0.52 ± 0.03*** | 1.17 ± 0.08 | 0.29 ± 0.02*** |
| Toc34 | 0.51 ± 0.03***<br>(0.44 ± 0.04)<sup>a</sup> | 0.98 ± 0.08<br>(0.81 ± 0.03)<sup>a</sup> | 0.11 ± 0.01**<br>(0.95 ± 0.07)<sup>a</sup> |
| Tic110 | 1.44 ± 0.11* | 0.88 ± 0.07 | 0.97 ± 0.05 |
| Tic40 | 1.24 ± 0.10 | 0.79 ± 0.06 | 0.68 ± 0.05 |
| Tic100 | 0.46 ± 0.05*** | 0.08 ± 0.02 | nd |
| Tic56 | 1.24 ± 0.11* | 0.12 ± 0.07 | 0.04 ± 0.01 |
| Tic20-I | 0.49 ± 0.05** | nd | nd |
| ClpC2 | 1.88 ± 0.12*** | 1.21 ± 0.11 | 1.45 ± 0.14 |
| ClpB3 | 1.09 ± 0.06 | 0.92 ± 0.08 | 1.12 ± 0.11 |
| cpHsc70-1 | 1.15 ± 0.07 | 1.17 ± 0.08 | 1.08 ± 0.09 |
| ptCpn60 | 0.82 ± 0.14 | 0.49 ± 0.06 | 0.33 ± 0.04* |

On the contrary, the subunit composition of the two chloroplast translocation complexes TOC and TIC, together with the stromal chaperone system showed few major differences between *gun1-102* and Col-0 seedlings grown on MS medium with and without lincomycin (Figure 4b and Table 1). On MS medium without lincomycin, Toc34 and Toc159, the GTP-dependent receptor subunits responsible for interactions with distinct regions of the N-terminal cTP (chloroplast transit peptide) of cytosolic pre-proteins, were decreased by about 40 and 50% in *gun1-102* cotyledons in comparison to Col-0, respectively. Furthermore, a Toc34-specific band of higher molecular weight was also detected in *gun1-102*. Interestingly, amounts of the protein-conducting channel Toc75 were also reduced by 50% in *gun1-102*, indicating that the entire heterotrimeric TOC complex is destabilized by the loss of GUN1 in Arabidopsis cotyledons. Conversely, the inner membrane channel Tic110 and the associated protein Tic40 showed a slight increase in the *gun1-102* sample, while the channel protein of the 1-MDa complex, Tic20-I, together with Tic100, were reduced to half their control levels. Besides the TIC and TOC complexes, the plastid translocation machinery requires the activity of various stromal chaperones, mainly the chloroplast cpHsc70, Hsp90, Hsp93 (also known as ClpC2) and ptCpn60, which are involved in folding newly imported proteins and/or providing the energy required for protein translocation, via ATP hydrolysis. However, only ClpC2 showed a two-fold increase in *gun1-102* chloroplasts, while the levels of ClpB3, cpHsc70-1 and ptCpn60 were almost unchanged from those in Col-0 cotyledons. However, differences at the level of the TIC-TOC translocation machinery were more pronounced in seedlings grown on MS medium with lincomycin. In particular, the heterotrimeric TOC complex was markedly destabilized in *gun1-102* + Lin cotyledons, since Toc75 was reduced by more than threefold in comparison to Col-0, and only about 10% of the Toc34 subunit was present in its mature form; the rest comprised four distinct bands of between 50 and 90 kDa. In addition, subunits of the 1-MDa complex, such as Tic20-I and Tic100, were below the limits of detection in *gun1-102* + Lin cotyledons and only a very faint band was observed for Tic100 in Col-0 + Lin samples. Similarly, the level of Tic56 was reduced to 12% of its normal amount in Col-0 + Lin and to 4% in *gun1-102* + Lin cotyledons. It can be argued that the lincomycin-mediated inhibition of Tic214 (YCF1) synthesis in the chloroplast prevents the stable accumulation of the entire 1-MDa TIC complex, although such an effect is more pronounced in *gun1-102* than in Col-0 cotyledons, most probably due to the reduced level of *Tic214* transcripts in *gun1-102* + Lin cotyledons (see Figure 1c). Indeed, total protein ubiquitination was markedly increased in Col-0 + Lin, and even more so in *gun1-102* + Lin cotyledons, with respect to the Col-0 and *gun1-102* samples, between which no major differences were observed (Figure 4c). These results again underline the greater sensitivity of *gun1-102* seedlings to lincomycin (Figure 2). Furthermore, immunoprecipitation of Toc34 subunit from Col-0 + Lin and *gun1-102* + Lin samples, followed by immunoblot analysis with the UBQ11-specific antibody, allowed us to clarify that the high molecular bands observed between 50 and 90 kDa are the result of Toc34 ubiquitination (Figure 4d).

### Plastid precursor proteins accumulate in mesophyll cells of *gun1* cotyledons

The alteration of the translocon machinery in cotyledons lacking the GUN1 protein prompted us to investigate if chloroplast-targeted proteins accumulated in the cytosol of mesophyll cells in *gun1-102* cotyledons. To this aim, the soluble fraction of total proteins extracted from cotyledons of Col-0 (+/- Lin) and *gun1-102* (+/- Lin) seedlings was analyzed by nLC-ESI-MS/MS. An equal amount of peptides, corresponding to 750 ng of proteins, was injected. Equal loading was confirmed by the total number of PSMs (Peptide-Spectrum Match) identified in each sample (Table S2). The peptides identified were then mapped to the amino-acid sequences of 1241 proteins listed in Uniprot (https://www.uniprot.org/proteomes/UP000006548) and known to have a cTP and on another 721 nucleus-encoded proteins predicted by ChloroP 1.1 (www.cbs.dtu.dk/services/ChloroP/) to be imported into the chloroplast. In Table 2, a list of chloroplast imported proteins for which at least one peptide was found in each of the three biological replicates of at least one of the four samples is reported. In total, a cTP was detected in 37 cotyledon proteins, of which 10 accumulated preferentially and with statistical significance in *gun1-102* + Lin samples (see bold ATG code in Table 2). Among these are subunits of the thylakoid electron transport chain and enzymes of the Calvin-Benson cycle. In general, a net and statistically significant increase in cTP PSMs was detected in *gun1-102* + Lin cotyledons, implying that the lack of GUN1, in combination with the presence of lincomycin in the growth medium, favors the accumulation of chloroplast precursor proteins in the cytosol.

**Table 2.**
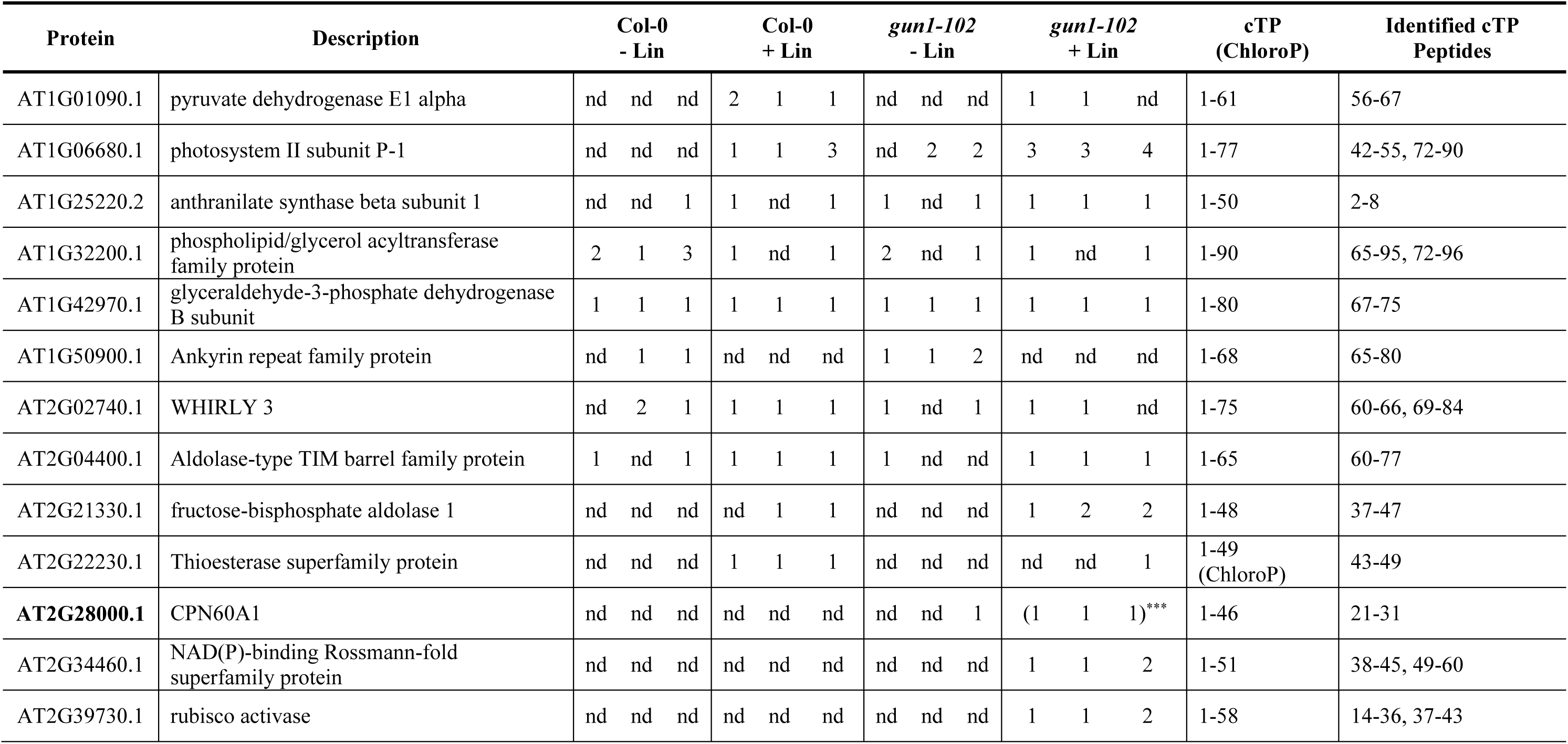
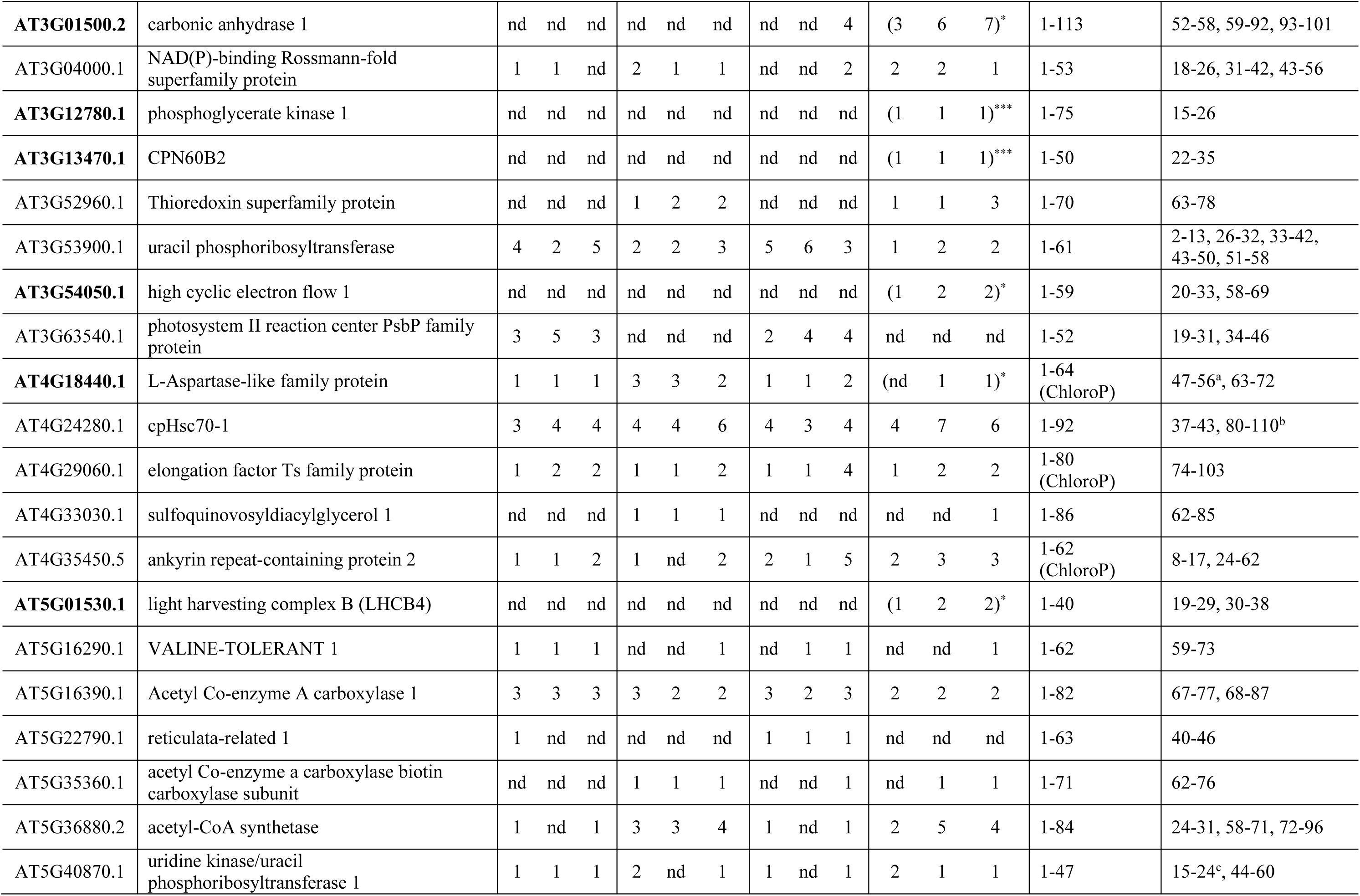
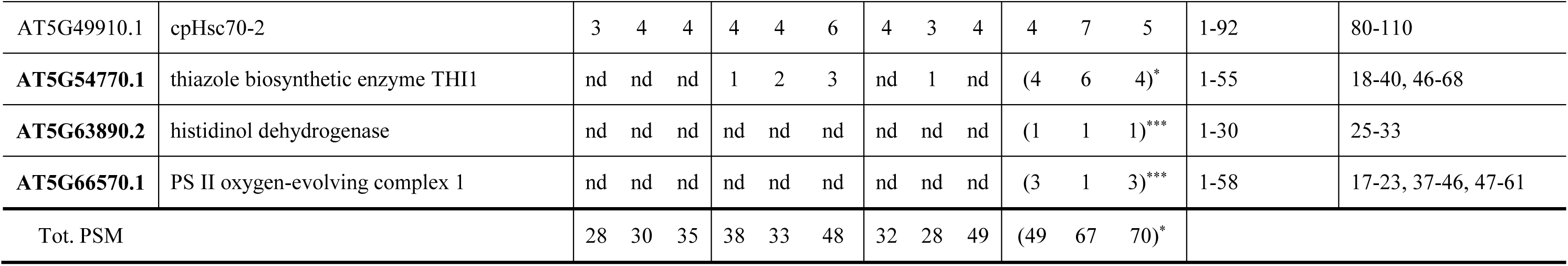
List of the nucleus-encoded precursor proteins targeted to the chloroplast detected in total cotyledon soluble extracts from Col-0 and gun1-102. At least one tryptic peptide derived from the chloroplast transit peptide (cTP) of each of the proteins listed was found in all three biological replicates of at least one of the four samples. The results are presented as the number of peptide spectrum matches (PSMs) corresponding to any cTP peptide found in each of the three biological replicates. The list of peptides identified in the total soluble extract has been manually screened for sequences falling within the cTP of one of the 1241 *Arabidopsis thaliana* entries with a known cTP available in Uniprot, or in one of the remaining 721 accessions annotated as targeted to the chloroplast. In the latter case, the sequences of the theoretical transit peptides were predicted using ChloroP 1.1 (www.cbs.dtu.dk/services/ChloroP/). cTP (ChoroP): sequence of the cTP as reported in Uniprot or, when marked (ChoroP), as predicted by ChloroP 1.1. Identified cTP peptides: starting- and ending-amino acid position of the sequences covered by the identified tryptic peptides in the cTP segment of the respective protein. Asterisks indicate the statistical significance with respect to Col-0 + Lin values, as evaluated by Student’s t-test and Welch correction (*, p<0.05; **, p<0.01; ***, p<0.001).

| Protein | Description | Col-0<br>- Lin | Col-0<br>+ Lin | *gun1-102*<br>- Lin | *gun1-102*<br>+ Lin | cTP<br>(ChloroP) | Identified cTP<br>Peptides |
| --- | --- | --- | --- | --- | --- | --- | --- |
| AT1G01090.1 | pyruvate dehydrogenase E1 alpha | nd nd nd | 2 1 1 | nd nd nd | 1 1 nd | 1-61 | 56-67 |
| AT1G06680.1 | photosystem II subunit P-1 | nd nd nd | 1 1 3 | nd 2 2 | 3 3 4 | 1-77 | 42-55, 72-90 |
| AT1G25220.2 | anthranilate synthase beta subunit 1 | nd nd 1 | 1 nd 1 | 1 nd 1 | 1 1 1 | 1-50 | 2-8 |
| AT1G32200.1 | phospholipid/glycerol acyltransferase family protein | 2 1 3 | 1 nd 1 | 2 nd 1 | 1 nd 1 | 1-90 | 65-95, 72-96 |
| AT1G42970.1 | glyceraldehyde-3-phosphate dehydrogenase B subunit | 1 1 1 | 1 1 1 | 1 1 1 | 1 1 1 | 1-80 | 67-75 |
| AT1G50900.1 | Ankyrin repeat family protein | nd 1 1 | nd nd nd | 1 1 2 | nd nd nd | 1-68 | 65-80 |
| AT2G02740.1 | WHIRLY 3 | nd 2 1 | 1 1 1 | 1 nd 1 | 1 1 nd | 1-75 | 60-66, 69-84 |
| AT2G04400.1 | Aldolase-type TIM barrel family protein | 1 nd 1 | 1 1 1 | 1 nd nd | 1 1 1 | 1-65 | 60-77 |
| AT2G21330.1 | fructose-bisphosphate aldolase 1 | nd nd nd | nd 1 1 | nd nd nd | 1 2 2 | 1-48 | 37-47 |
| AT2G22230.1 | Thioesterase superfamily protein | nd nd nd | 1 1 1 | nd nd nd | nd nd 1 | 1-49<br>(ChloroP) | 43-49 |
| **AT2G28000.1** | CPN60A1 | nd nd nd | nd nd nd | nd nd 1 | (1 1 1)<sup>***</sup> | 1-46 | 21-31 |
| AT2G34460.1 | NAD(P)-binding Rossmann-fold superfamily protein | nd nd nd | nd nd nd | nd nd nd | 1 1 2 | 1-51 | 38-45, 49-60 |
| AT2G39730.1 | rubisco activase | nd nd nd | nd nd nd | nd nd nd | 1 1 2 | 1-58 | 14-36, 37-43 |
| **AT3G01500.2** | carbonic anhydrase 1 | nd nd nd | nd nd nd | nd nd 4 | (3 6 7)* | 1-113 | 52-58, 59-92, 93-101 |
| AT3G04000.1 | NAD(P)-binding Rossmann-fold superfamily protein | 1 1 nd | 2 1 1 | nd nd 2 | 2 2 1 | 1-53 | 18-26, 31-42, 43-56 |
| **AT3G12780.1** | phosphoglycerate kinase 1 | nd nd nd | nd nd nd | nd nd nd | (1 1 1)*** | 1-75 | 15-26 |
| **AT3G13470.1** | CPN60B2 | nd nd nd | nd nd nd | nd nd nd | (1 1 1)*** | 1-50 | 22-35 |
| AT3G52960.1 | Thioredoxin superfamily protein | nd nd nd | 1 2 2 | nd nd nd | 1 1 3 | 1-70 | 63-78 |
| AT3G53900.1 | uracil phosphoribosyltransferase | 4 2 5 | 2 2 3 | 5 6 3 | 1 2 2 | 1-61 | 2-13, 26-32, 33-42, 43-50, 51-58 |
| **AT3G54050.1** | high cyclic electron flow 1 | nd nd nd | nd nd nd | nd nd nd | (1 2 2)* | 1-59 | 20-33, 58-69 |
| AT3G63540.1 | photosystem II reaction center PsbP family protein | 3 5 3 | nd nd nd | 2 4 4 | nd nd nd | 1-52 | 19-31, 34-46 |
| **AT4G18440.1** | L-Aspartase-like family protein | 1 1 1 | 3 3 2 | 1 1 2 | (nd 1 1)* | 1-64<br>(ChloroP) | 47-56<sup>a</sup>, 63-72 |
| AT4G24280.1 | cpHsc70-1 | 3 4 4 | 4 4 6 | 4 3 4 | 4 7 6 | 1-92 | 37-43, 80-110<sup>b</sup> |
| AT4G29060.1 | elongation factor Ts family protein | 1 2 2 | 1 1 2 | 1 1 4 | 1 2 2 | 1-80<br>(ChloroP) | 74-103 |
| AT4G33030.1 | sulfoquinovosyldiacylglycerol 1 | nd nd nd | 1 1 1 | nd nd nd | nd nd 1 | 1-86 | 62-85 |
| AT4G35450.5 | ankyrin repeat-containing protein 2 | 1 1 2 | 1 nd 2 | 2 1 5 | 2 3 3 | 1-62<br>(ChloroP) | 8-17, 24-62 |
| **AT5G01530.1** | light harvesting complex B (LHCB4) | nd nd nd | nd nd nd | nd nd nd | (1 2 2)* | 1-40 | 19-29, 30-38 |
| AT5G16290.1 | VALINE-TOLERANT 1 | 1 1 1 | nd nd 1 | nd 1 1 | nd nd 1 | 1-62 | 59-73 |
| AT5G16390.1 | Acetyl Co-enzyme A carboxylase 1 | 3 3 3 | 3 2 2 | 3 2 3 | 2 2 2 | 1-82 | 67-77, 68-87 |
| AT5G22790.1 | reticulata-related 1 | 1 nd nd | nd nd nd | 1 1 1 | nd nd nd | 1-63 | 40-46 |
| AT5G35360.1 | acetyl Co-enzyme a carboxylase biotin carboxylase subunit | nd nd nd | 1 1 1 | nd nd 1 | nd 1 1 | 1-71 | 62-76 |
| AT5G36880.2 | acetyl-CoA synthetase | 1 nd 1 | 3 3 4 | 1 nd 1 | 2 5 4 | 1-84 | 24-31, 58-71, 72-96 |
| AT5G40870.1 | uridine kinase/uracil phosphoribosyltransferase 1 | 1 1 1 | 2 nd 1 | 1 nd 1 | 2 1 1 | 1-47 | 15-24<sup>c</sup>, 44-60 |
| AT5G49910.1 | cpHsc70-2 | 3 4 4 | 4 4 6 | 4 3 4 | 4 7 5 | 1-92 | 80-110 |
| **AT5G54770.1** | thiazole biosynthetic enzyme THI1 | nd nd nd | 1 2 3 | nd 1 nd | (4 6 4)* | 1-55 | 18-40, 46-68 |
| **AT5G63890.2** | histidinol dehydrogenase | nd nd nd | nd nd nd | nd nd nd | (1 1 1)*** | 1-30 | 25-33 |
| **AT5G66570.1** | PS II oxygen-evolving complex 1 | nd nd nd | nd nd nd | nd nd nd | (3 1 3)*** | 1-58 | 17-23, 37-46, 47-61 |
| Tot. PSM |  | 28 30 35 | 38 33 48 | 32 28 49 | (49 67 70)* |  |  |

The existence of a defect in the import of chloroplast proteins in *gun1-102* cotyledons was verified further by immunoblot analyses on total protein extracts from Col-0 and *gun1-102* seedlings grown on MS medium in the presence of lincomycin, using FtsH1, FtsH2 and FtsH5 antibodies (Figure 5a). A marked reduction in FtsH proteins was observed in cotyledons of Lin-grown seedlings, together with the appearance of FtsH-specific bands of higher molecular weight (hwFtsH) migrating between 70 and 80 kDa, which are compatible with the apparent molecular weights of FtsH precursors. Interestingly, the hwFtsH forms were considerably more abundant in Lin-grown *gun1-102* cotyledons, but they were also detectable in *gun1-102* seedlings grown on standard MS medium, when FtsH1-, FtsH2- and FtsH5-specific antibodies were employed for immunodetection (Figure 5a). The hwFtsH proteins could be fully recovered in the extra-plastidial (Sup) soluble fraction of cotyledon proteins after chloroplasts were isolated from lincomycin-grown seedlings, suggesting that they could represent FtsH1, FtsH2 and FtsH5 proteins that retained their chloroplast and thylakoid (tTP) transit peptides (Figure 5b). To avoid problems due to the cross-reactivity of FtsH antibodies, the coding sequences of *FtsH1*, *FtsH2, FtsH5* and *FtsH8* genes were placed under the control of the CaMV 35S promoter and fused in frame to either *GFP* (*oeFtsH1-GFP*, *oeFtsH2-GFP* and *oeFtsH8-GFP*) or *RFP* (*oeFtsH5-RFP*). These constructs were introduced, via Agrobacterium-mediated transformation, into Col-0 and the corresponding *gun1-102 ftsh* double mutants, and the accumulation pattern of the chimeric proteins was monitored in control (non-transformed Col-0 seedlings) and transformed seedlings grown on MS medium with and without lincomycin (Figure 5c). In agreement with the observations made on the endogenous FtsH proteins (see Figure 5a), using GFP- and RFP-specific antibodies as appropriate, higher-molecular-weight proteins were detected in *gun1-102 ftsh* mutant cotyledons grown on lincomycin-containing medium and expressing FtsH1-GFP (*gun1-102 ftsh1-1* + *oeFtsH1-GFP*), FtsH2-GFP (*gun1-9 ftsh2-3 + oeFtsH2-GFP*), or FtsH5-RFP (*gun1-102 ftsh5-3 + oeFtsH5-RFP*). In addition, high-molecular-weight FtsH2-GFP- and FtsH5-RFP-specific signals were also detected in *gun1-9 ftsh2-3 + oeFtsH2-GFP* and *gun1-102 ftsh5-3 + oeFtsH5-RFP* genetic backgrounds grown in the absence of the plastid translation inhibitor, further supporting a role for GUN1 in plastid protein import.

**Figure 5.**
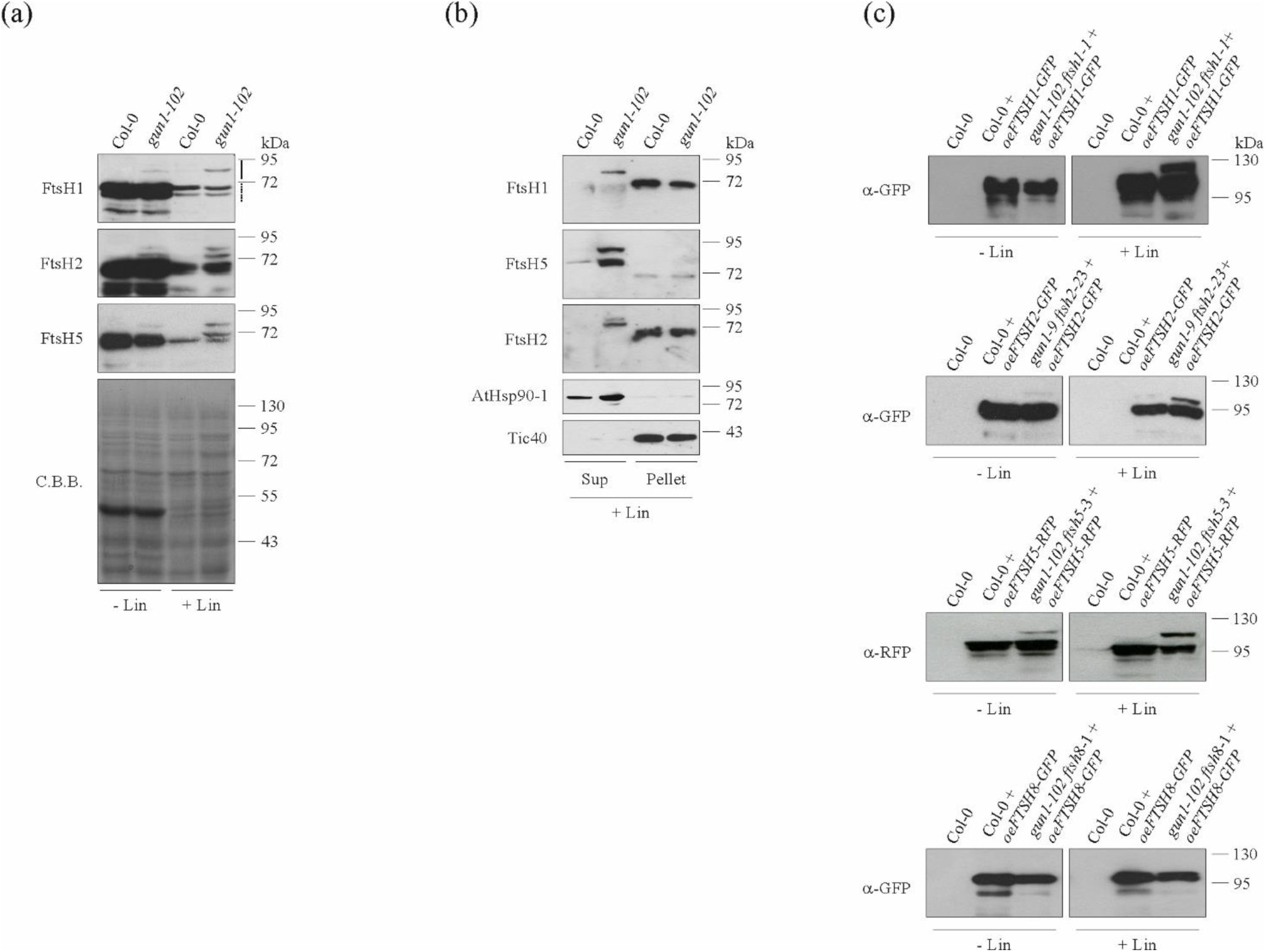
Accumulation of FtsH precursors in Col-0 and *gun1-102* seedlings grown on MS medium with and without lincomycin. **(a)** Immunoblot analyses of total proteins extracted from Col-0 (+/- Lin) and *gun1-102* (+/- Lin) cotyledons. Filters were immunodecorated with antibodies specific for the FtsH1, FtsH2, FtsH5 proteins and the small subunit of RUBISCO (RbcS). A Coomassie Brilliant Blue (C.B.B.)-stained gel, prominently displaying the RUBISCO large subunit (RbcL), is shown as loading control. The continuous line to the right of the FtsH1 immunoblot indicates the hwFtsH region, while the dotted line shows the mFtsH segment. **(b)** The soluble, extra-chloroplast fraction (Sup) and the plastid-enriched fraction (Pellet) were isolated from Col-0 and *gun1-102* cotyledons obtained from seedlings grown on MS medium + Lin. After SDS-PAGE fractionation and blotting, filters were immunolabeled with FtsH1-, FtsH2- and FtsH5-specific antibodies. The AtHsp90-1 antibody was used as a marker for the soluble fraction, while Tic40 was used for the plastid-enriched fraction. **(c)** Immunoblots of total protein extracts from cotyledons of non-transformed Col-0 seedlings, as well as of Col-0 and different *gun1 ftsh* double mutant seedlings transformed with *35S::FtsH-GFP/RFP* (*oeFtsH1/2/8-GFP* and *oeFtsH5-RFP*) constructs, and grown on MS medium with and without Lin. Commercially available GFP- and RFP-specific antibodies were used to detect the FtsH chimeras. Five mg (fresh weight) samples of cotyledon tissue, collected at 6 DAS, were used for the analyses.

The accumulation of plastid precursor proteins was also verified by nLC-ESI-MS/MS analysis of trypsin-digested slices isolated from the 75- to 90-kDa and the 60-to-70 kDa regions of gel lanes loaded with Col-0 (+/- Lin) and *gun1-102* (+/-Lin) extracts, indicated as hwFtsH bands and as mature protein bands (mFtsH) in Figure 5a. The proteins identified in the four hwFtsH and the four mFtsH bands are listed in Table S3. The four subunits of the thylakoid FtsH protease complex were identified with high confidence in the mFtsH bands of all four samples, and the number of PSMs corresponding to peptides unique to one or common to all four FtsH subunits were considerably more abundant in both Col-0 and *gun1-102* lanes than in Col-0 + Lin and *gun1-102* + Lin samples, in agreement with the marked reduction in the levels of thylakoid FtsH complexes in seedlings grown in the presence of lincomycin, as shown by immunoblot analysis (see Figure 5a). However, only the FtsH1 subunit was found with a high confidence in the hwFtsH band from *gun1-102* + Lin cotyledons (Table S3). To increase the chances of detecting FtsH2 and FtsH5 peptides, all eight samples (four hwFtsH and four mFtsH bands) were re-analyzed by monitoring only the masses corresponding to tryptic peptides unique to FtsH2 and FtsH5 (as predicted by *in-silico* digestion of their complete sequences; see Table S4) in both mature and precursor bands (Tables S5 and S6). In this case, peptides of the mature FtsH5 were identified in all mature and all precursor bands (Table S6). In addition, the precursor band in the *gun1-102* + Lin sample contained two peptides derived from the FtsH5-cTP: [K].SLPFSVISR.[K] and [R].YQISQSEK.[L], and the FtsH5-cTP peptide [K].SLPFSVISR.[K] was also detected in the precursor band in the *gun1-102* sample. FtsH2 peptides were primarily found in the mFtsH bands of the four samples (Table S5). No FtsH2-cTP peptide was detected among the eight bands analyzed, although the [K].ILLGNAGVGLVASGK.[A] peptide – part of the thylakoid transit peptide – was found in the mature band of *gun1-102* + Lin.

Furthermore, the accumulation of precursor proteins was corroborated by the increased accumulation of the cytosolic Heat Shock Protein 90 (AtHsp90) and the Heat Shock cognate protein 70 (AtHsc70) in *gun1-102* + Lin cotyledons, detected by trypsin digestion of gel slices (see asterisks in Figure 6a) loaded with Col-0 + Lin or *gun1-102* + Lin samples, followed by liquid chromatography-mass spectrometry (LC-ESI MS/MS) analyses (Figure 6b and Table S7). Moreover, immunoblot analysis of AtHsp90-1 and AtHsc70-4 using specific antibodies confirmed the increased amounts of heat-shock proteins in the cytosol of *gun1-102* cotyledon cells grown in the presence of lincomycin, (Figure 6c). Indeed, members of Hsp90 protein family are reported to assist the transport of precursor proteins into organelles or, as in the case of Hsc70-4, to be involved in the degradation of chloroplast pre-proteins that accumulate in the cytosol.

**Figure 6.**
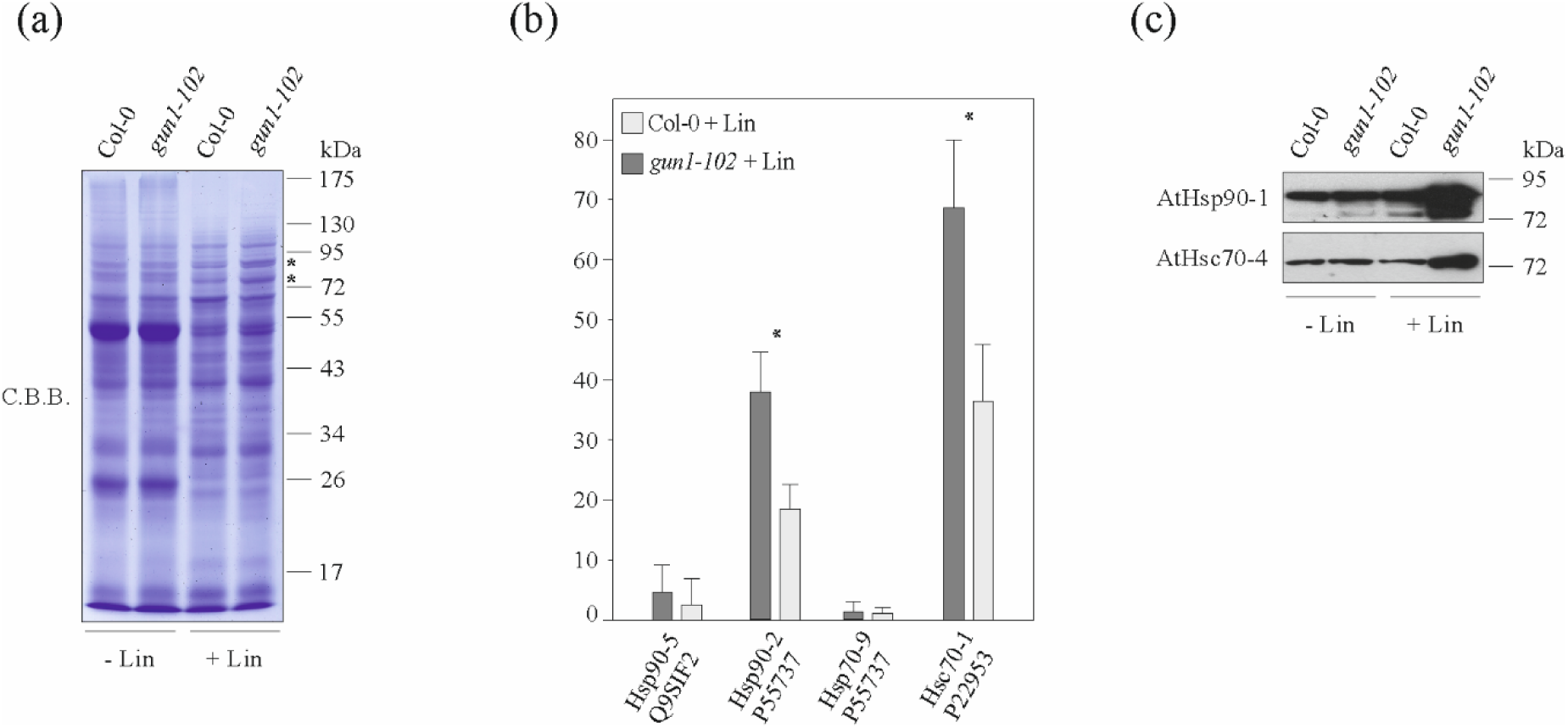
Liquid chromatography-mass spectrometry (LC-ESI MS/MS) analysis of proteins migrating in the 75- to 90-kDa region of SDS-PA gels. **(a)** C.B.B. stained SDS-PA gel loaded with total protein extracts from Col-0 (+/- Lin) and *gun1-102* (+/-) cotyledons, which was used to isolate the protein bands at ∼ 75 kDa and ∼ 90 kDa (see asterisks) that are highly abundant in *gun1-102* + Lin seedlings. **(b)** The gel slices indicated by the asterisks were incubated with trypsin, and the resulting peptides were analyzed by LC-ESI MS/MS. The chart shows the relative levels (expressed as number of peptides/number of total peptides per gel slice) of four detected HSP protein families in Col-0 + Lin and *gun1-102* + Lin samples. The family name was assigned based on the top rank hits. Mean values ± SD are provided. Asterisks indicate the statistical significance, as evaluated by Student’s t-test (p<0.05). **(c)** Immunoblots of total protein extracts from Col-0 (+/- Lin) and *gun1-102* (+/- Lin) cotyledons (see SDS-PAGE in panel a) probed with antibodies specific for the two cytosolic chaperones AtHsc70-4 and AtHsp90-1. Images of filters representative of three biological replicates are shown.

### The activities of GUN1 and proteins involved in chloroplast protein homeostasis are essential for correct chloroplast biogenesis in cotyledons

To further investigate the importance of GUN1 activity in chloroplast biogenesis upon alteration of plastid protein homeostasis (see also Tadini et. al., 2016 and Paieri et al., 2018), the *gun1 ftsh*, *gun1-102 cphsc70-1* and *gun1-102 sca3-1* double mutant seedlings were also analyzed at macroscopic level (Figure 7a and Figure S3a). Specifically, mutations in the nuclear genes for FtsH1 (*ftsh1-1*), FtsH5 (*ftsh5-3*) and FtsH8 (*ftsh8-1*) proteins were introduced into the *gun1-102* mutant background by manual crossing, while the CRISPR-Cas9 editing strategy was used to generate the *ftsh2-3* mutant allele in the *gun1-9* genetic background, since these two genes are closely linked on chromosome 2 (Figure S1). Notably, *gun1-102, gun1-9*, *ftsh1-1, ftsh5-3* and *ftsh8-1* single mutants displayed wild-type-like cotyledons in terms of chlorophyll content (Figure 7b) and photosynthetic activity (Figure 7c), while *ftsh2-2* seedlings showed a slight reduction in chlorophyll content and photosynthetic performance. Furthermore, while *gun1-102 ftsh1-1* and *gun1-102 ftsh8-1* homozygotes were indistinguishable from Col-0 seedlings (Figure S3a), the double mutants *gun1-102 ftsh5-3* and *gun1-9 ftsh2-3* displayed exacerbated phenotypes characterized by severely variegated cotyledons, i.e. reduced chlorophyll content and low photosynthesis efficiency (Figure 7). The enhanced variegated phenotype of the *gun1-102 ftsh5-3* double mutant was observable up to the four-leaf rosette stage, but at later stages (21 DAS) these mutants appeared identical to *ftsh5* plants. In contrast, the *gun1-9 ftsh2-3* double-mutant phenotype resulted in lethality at the two-leaf rosette stage. To obtain a more general view of the importance of GUN1 for plastid protein homeostasis, the *gun1-102* mutation was also introduced into the *cphsc70-1* mutant background, which lacks the chloroplast heat shock protein 70-6 (Figure 7a). Indeed, the *gun1-102 cphsc70-1* seedlings showed altered greening of cotyledons, with the appearance of white sectors typical of the variegated phenotype, confirming further the role of GUN1 in chloroplast biogenesis upon perturbation of chloroplast protein homeostasis in Arabidopsis cotyledons. Similarly, the absence of GUN1 protein together with a drastic reduction of NEP enzyme, as in *gun1-102 sca3-1*, resulted in albino seedlings. Furthermore, it is interesting to observe that the *sca3-1* single mutant, characterized by pale-green cotyledons (Figure 7a) shows an increased sensitivity to lincomycin, even higher than *gun1-102*, as can be observe by the marked drop of PSII yield already at 5.5 µM of the plastid translation inhibitor (Figure 2), most probably as a consequence of the complete inhibition of the Δ-*rpo* compensatory response, similarly to the case of *gun1-102* seedlings (see *rps12-3’* accumulation pattern in Figure S3b). Nevertheless, the *sca3-1* mutant does not show a gun phenotype as can be observed by the marked drop of *rbcs* transcripts upon lincomycin treatment (Figure S3b).

**Figure 7.**
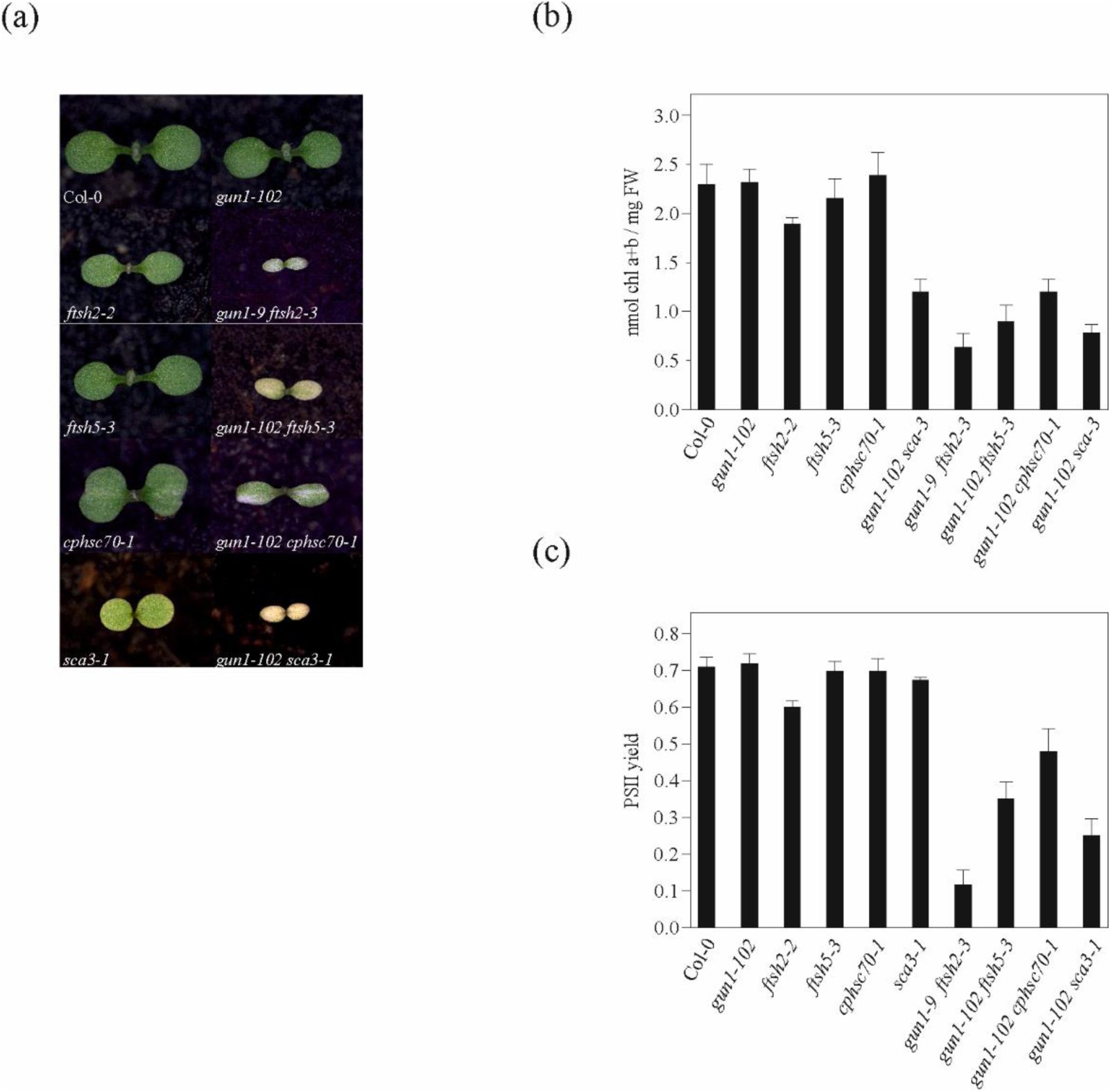
Genetic interactions between GUN1, subunits of the thylakoid FtsH protease complex and the chloroplast cpHsc70 protein. **(a)** Phenotypes of single *gun1-102*, *ftsh*, *cphsc70-1*, *sca3-1* mutants and double mutants at 6 days after sowing (DAS), when only the two cotyledons have yet been formed. **(b)** Cotyledon chlorophyll content of lines shown in (a), measured at 6 DAS. The total chlorophyll content is expressed relative to cotyledon fresh weight (nmol chl a+b /mg FW). **(c)** The photosynthetic performance of cotyledons of the lines shown in (a) was measured as the effective quantum yield of PSII, employing a DUAL-PAM-100 (WALZ). Note that *gun1-9* showed the same visible phenotype, chlorophyll content and photosynthetic performance as *gun1-102* seedlings.

To better understand the cotyledon phenotypes of mutant seedlings, we analyzed the ultrastructure of mesophyll cell chloroplasts by transmission electron microscopy (TEM; Figure 8). TEM of thin sections of cotyledons from Col-0, *gun1-102* and *ftsh5-3* seedlings (grown at 100 μmol photons m-2 sec-1) showed the characteristic chloroplast organization, including stacked grana thylakoids, stroma lamellae and starch granules. In contrast, the paler cotyledons of *ftsh2-2* seedlings were characterized by highly vacuolated, rounded chloroplasts lacking the typical thylakoid organization in grana and stroma lamellae. In addition, vesicle-like structures were observed at the border between the chloroplast and the tonoplast in the case of *ftsh2-2* chloroplasts. Introduction of the *gun1-102* mutation into the *ftsh5-3* background led to the appearance of several cotyledon mesophyll cells in which fully developed wild-type-like chloroplasts were replaced by plastids devoid of thylakoid membranes and with large vesicles budding from the organellar envelope, which were delivered to the vacuole. A more drastic cotyledon phenotype was seen in *gun1-9 ftsh2-3* seedlings, where highly vacuolated, irregularly shaped chloroplasts were rarely observed in mesophyll cells, and a considerable number of highly electron-dense structures – possibly the products of plastid degradation – were observed within vacuoles (Figure 8). Overall, it can be argued that the lack of the Δ-*rpo* compensatory response, together with the plastid-to-nucleus communication, in *gun1* cotyledons add to the alteration of chloroplast protein homeostasis in *ftsh* and *cphsc70-1* and *sca3-1* mutants (see Figure 1, Figure 2, Figure S2, Figure S3 and Table S1), impairing chloroplast biogenesis as observed previously (Tadini et al., 2016 and Paieri et al., 2018).

**Figure 8.**
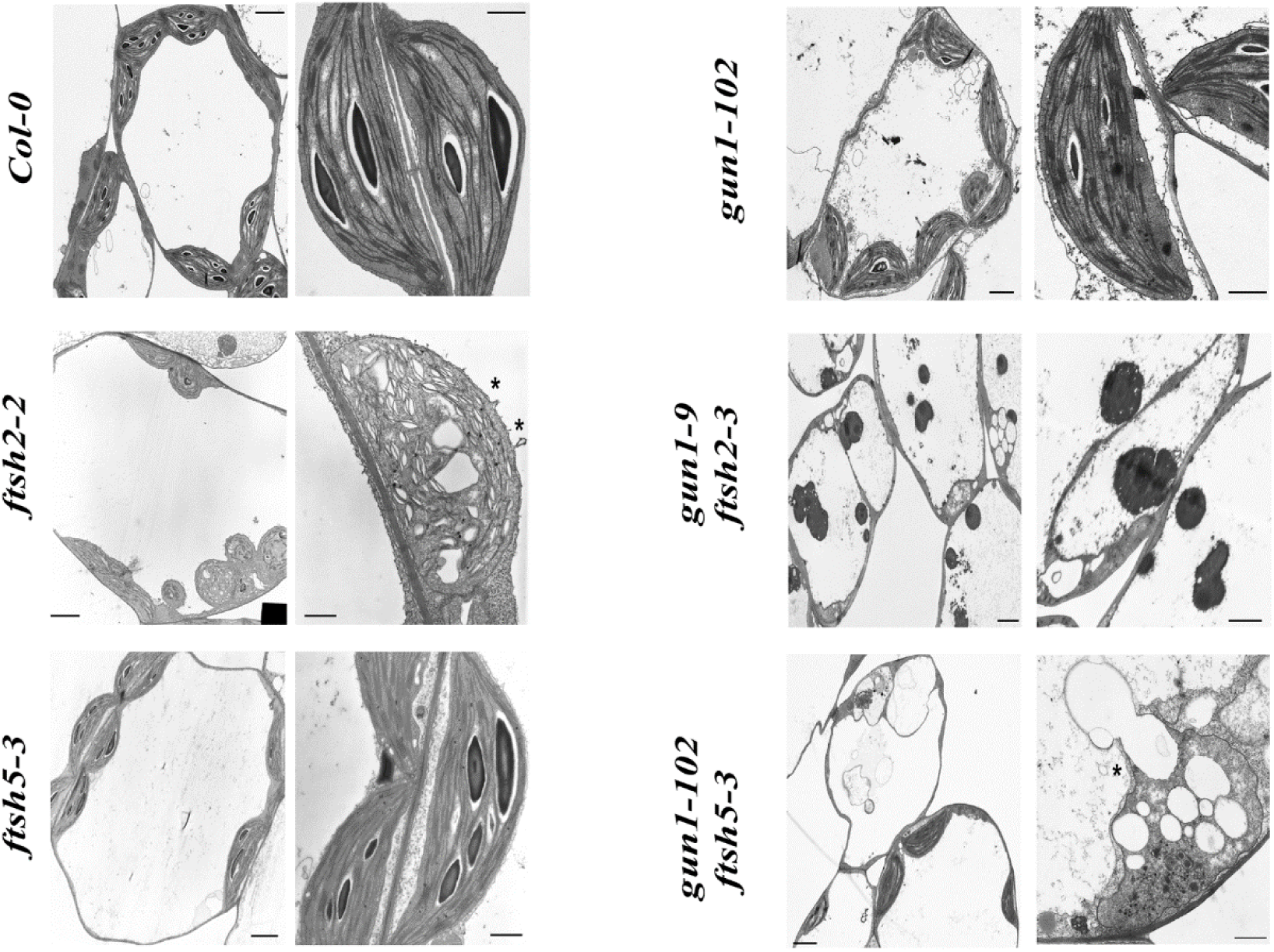
TEM micrographs of chloroplasts in mesophyll cells of 6-day-old Arabidopsis cotyledons from Col-0, and *gun1-102*, *ftsh5-3*, *ftsh2-2* single mutants and *gun1-102 ftsh5-3* and *gun1-9 ftsh2-3* double mutants. Ultrathin sections of cotyledons from Col-0 and mutant seedlings were stained with 2% uranyl acetate and lead citrate and examined by TEM. Scale bars correspond to 5 μm and 1 μm. Note that the WT-like ultrastructure of *gun1-102* chloroplasts was also observed in *gun1-9* cotyledons. Asterisks (*) indicate vesicles budding from the chloroplast envelope.

## Discussion

The major goal of this study was to clarify the role of GUN1 during early stages of chloroplast biogenesis in Arabidopsis cotyledons, as previously done in mature rosette leaves (Tadini et al., 2016). A major difficulty in the elucidation of the molecular mechanism and the precise role of GUN1 in retrograde signaling arises from the fact that *gun1* mutants are virtually indistinguishable from wild type, with only a small percentage of *gun1* mutants developing chlorophyll-deficient cotyledons under optimal growth conditions (Ruckle et al., 2007). However, seedlings lacking GUN1 are hypersensitive to low concentrations of both norflurazon (20 nM) and lincomycin (22 µM), as revealed by a small pale yellow or white cotyledon phenotype, while wild-type seedlings form fully developed green cotyledons under the same growth conditions (Song et al., 2018; Zhao et al., 2018; see also Figure S2a). Accordingly, when the *gun1-102* mutant was combined with mutations that impair plastid protein synthesis, such as *prps17-1*, *prpl11-1* and *prpl24-1*, the corresponding double mutants were seedling lethal (Tadini et al., 2016; Paieri et al., 2018). A similar phenotype was observed in double mutants defective in both GUN1 and Clp protease activity, supporting the notion that GUN1 is essential for protein homeostasis in the chloroplast (Llamas et al., 2017). Our data further support these inferences, as shown by the seedling lethal phenotypes of *gun1-9 ftsh2-3* and *gun1-102 sca3-1* and the enhanced variegation phenotype of *gun1-102 ftsh5-3* and *gun1-102 cphsc70-1* cotyledons. This notion is also corroborated by the differences in chloroplast ultrastructure associated with depletion of FtsH alone or together with GUN1, as in *gun1-102 ftsh5-3* and *gun1-9 ftsh2-3* seedlings (see also Figure 7a and 8).

### GUN1 plays a key role in the NEP-dependent compensatory response

Since GUN1 belongs to the large PPR protein family whose members are involved in different aspects of RNA metabolism, we have explored the role of GUN1 in plastid transcript accumulation upon perturbation of plastid protein homeostasis in Arabidopsis Col-0 and mutant cotyledons. In particular, the addition of lincomycin to the growth medium abolished the accumulation of PEP-dependent transcripts and dramatically increased that of NEP-dependent mRNAs, both in Col-0 and mutant cotyledons. Strikingly, cotyledons lacking GUN1 showed very limited changes in the abundance of NEP-dependent transcripts upon partial or total inhibition of PEP activity, as in the case of *sca3-1* mutant where the abundance of RpoT enzyme is markedly reduced. These findings together with the increased accumulation of NEP-dependent transcripts upon incubation of recGUN1 protein with *gun1-102* plastids in the Run on assay, support the notion that GUN1 is required to enhance NEP-dependent transcription in response to plastid protein homeostasis perturbation. The molecular mechanisms responsible for this abnormal transcription pattern, referred to as the “Δ-*rpo* phenotype”, are largely unknown. However, PLASTID REDOX INSENSITIVE2 (PRIN2), a redox-regulated protein, has recently been proposed as the link between photosynthetic electron transport and activation of PEP-dependent photosynthetic gene expression (Díaz et al., 2018). The physical interaction of GUN1 with NEP, as revealed by co-immunoprecipitation studies and BiFC assay (see Figure 3), could be one of the key elements of GUN1-mediated plastid transcript accumulation. Interestingly, homologs of GUN1 and PRIN2 are absent from the genomes of green algae and cyanobacteria, indicating that these proteins appeared concomitantly with the emergence of the nucleus-encoded plastid RNA polymerase NEP during the evolution of the plant lineage.

### Regulation of the translocon machinery is an important element of plastid protein homeostasis

The effects of perturbation of plastid protein homeostasis are not limited to chloroplast transcript accumulation and thylakoid membrane ultrastructure, but also impinge on the accumulation of nucleus-encoded proteins either located in the cytosol or targeted to the chloroplast and involved in different plastid activities. In addition, major changes in protein accumulation and post-translation modification were also observed in the envelope – in particular with respect to subunits of the translocon machinery (Figure 4b and Table 1). In general, loss of GUN1 destabilizes the entire TOC complex in Arabidopsis cotyledons, as shown by the reduced abundance of the GTP-dependent receptor protein Toc159, and the protein conducting channel Toc75. Furthermore, the other receptor protein Toc34 has been shown here to be ubiquitinated in *gun1-102* cotyledons, supporting previous reports that ubiquitination of TOC components serves to adjust the plastid proteome to developmental and physiological needs (Svozil et al., 2014). Similarly, the reduced accumulation of Tic100 and Tic20-I indicates that GUN1 is important for the accumulation/maintenance of the 1-MDa TIC complex. Notably, the presence of lincomycin in the growth medium further exacerbated the effects on the translocon machinery in terms of protein abundance and Toc34 ubiquitination. For instance, levels of the Tic20-I protein fall in the presence of lincomycin, in agreement with previous reports (Bolter et al., 1998; Köhler et al., 2016), and steady-state amounts of Tic100 and Tic56 are drastically reduced, which suggests the existence of a regulatory mechanism that coordinates chloroplast protein synthesis with chloroplast protein import. This postulate could also explain why the NEP-dependent *Tic214* (*Ycf1*) gene has been retained in the plastid genome of diverse plant species. In contrast, levels of Tic110 and Tic40 increased in *gun1-102* cotyledons and remained relatively high even in the presence of lincomycin, in agreement with the fact that Tic110 has been reported to be a target of the Clp protease in Chlamydomonas (Ramundo et al., 2014).

### Retrograde signaling and the ubiquitin-proteasome system

As described above, the chloroplast’s capacity for protein import is altered in *gun1-102* and even more so in *gun1-102* seedlings grown in the presence of lincomycin. As a consequence, chloroplast precursor proteins accumulate particularly in the cytosol of *gun1-102* cells. This accumulation of precursor proteins is compatible with the large increases in levels of the chaperones AtHsp90 and AtHsc70 and the marked rise in ubiquitinated proteins observed in *gun1-102* cotyledons grown on lincomycin. Interestingly, the cytosolic heat shock cognate Hsc70-4 and its interacting E3 ubiquitin ligase CHIP (Lee et al., 2009) mediate the degradation of unimported chloroplast precursor proteins by the ubiquitin-proteasome system, indicating that under conditions in which processes within the chloroplast are compromised, cytoplasmic protein degradation is enhanced. Hence, the degradation of unimported proteins via the ubiquitin-proteasome system probably plays a key role in mediating changes in the composition of the chloroplast proteome in response to intracellular or environmental signals. At least two other mechanisms involving the ubiquitin-proteasome pathway have been shown to play a role in retrograde signaling (Jarvis and López-Juez, 2013; Woodson, 2016). One is based on direct regulation of the protein translocation machinery. In this mechanism, SP1, a RING-type ubiquitin ligase located in the outer chloroplast membrane directly regulates the content of the TOC complex. In addition, ubiquitin-proteasome-dependent post-translational regulation has been also shown to modulate levels of the transcription factor GOLDEN2-LIKE 1 (GLK1), which is known to positively regulate the expression of photosynthesis-related genes during chloroplast biogenesis (Tokumaru et al., 2017). Thus our results, together with previous reports, indicate that ubiquitination and the 26S proteasome are integral parts of retrograde signaling pathways, together with the reprogramming of nuclear transcription, and all are engaged in the maintenance of functional chloroplasts within plant cells.

### Several protein interactors for a very low abundant protein

During the reviewing process of this manuscript, a new paper from the lab of Ralph Bock (Wu et al. 2019) was published, clearly supporting the role of GUN1 in chloroplast protein import and demonstrating that overaccumulation of unimported precursor proteins in the cytosol plays a role in retrograde signaling. Although our findings largely overlap with Bock’s data, the primary function of GUN1 inside the chloroplasts of Arabidopsis cotyledons, i.e. a direct involvement in protein import through interaction with the import-related chaperone cpHSC70-1 (Wu et al. 2019), rather than its direct role in the accumulation of NEP-dependent transcripts and, indirectly, in the efficiency of the chloroplast protein import machinery (see Figure 1, 3 and 4, this manuscript), remains controversial. Certainly, the exacerbated phenotypes observed in *gun1-102 cphsc70-1*, and *gun1-102 sca3-1* seedlings support multiple roles of GUN1 protein in maintenance of chloroplast protein homeostasis. Nevertheless, it still remains to be clarified how GUN1, a very low abundant protein, can control the activity and/or modulate the formation of protein complexes among highly abundant proteins (Wu et al., 2019; Tadini et al., 2016). On the contrary RpoTp (NEP), as GUN1, belongs to the very few plant proteins never identified in plant proteomics studies, indicating that they are both very low abundant and pointing to a RpoTp-GUN1 protein complex with a 1-to-1 subunit stoichiometric ratio, essential to favor the transcription of NEP-dependent genes upon depletion of PEP activity. This is also in line with the transient accumulation of RpoTp and GUN1 proteins during early stages of chloroplast biogenesis. Furthermore, a direct role of GUN1 in RNA editing by physically interacting with the MULTIPLE ORGANELLAR RNA EDITING FACTOR 2 (MORF2), a member of the so-called plastid RNA editosome has been reported, recently (Zhao et al., 2019). In particular, it was shown that the *gun1* mutant cotyledons have differential efficiency of RNA editing (C->U) levels of 11 sites in the plastid transcriptome, after norflurazon or lincomycin treatment, compared to WT. Intriguingly, the target transcripts were NEP-dependent, including RNAs of PEP core subunits β and β’ (Zhao et al., 2019). In this scenario, GUN1 appears to be part of a protein complex that acts as a positive direct regulator of NEP activity and, indirectly, of PEP-dependent transcription. Such protein complex could contribute to the increase of cellular RNA amount observed during germination or even to the doubling of plant cell RNA detected within 48 h under stress conditions, as previously reported (Zhang et al., 2011).

## Experimental Procedures

### Plant Material and Growth Conditions

The Arabidopsis (*Arabidopsis thaliana*, genetic background Col-0) T-DNA insertion mutant lines used in this work are listed in Table S8, together with *gun1-9* and *scabra3-1* mutant alleles caused by point mutations. In order to determine the insertion sites, the regions flanking the T-DNA insertions (Figure S1) were PCR-amplified and sequenced (primer sequences in Table S8). Multiple mutants were generated by manual crossing and PCR-based segregation, with the exception of *gun1-9 ftsh2-3*, which required a CRISPR-Cas9 genome editing strategy. *gun1-9 ftsh2-3* was generated by targeting the first exon of the *FtsH2* locus in the *gun1-9* mutant background, using the pDe-CAS9 vector described by Fauser et al., (2014). Mutant plants, carrying the mutation of interest and devoid of the CRISPR-Cas9 gene, were selected in the T3 generation. Primer sequences used for guide RNA design are listed in Table S8. The transgenic lines *oeFtsH1-GFP*, *oeFtsH2*-*GFP*, *oeFtsH5-RFP* and *oeFtsH8-GFP* were generated by introducing *FtsH1*, *FtsH2*, *FtsH5* and *FtsH8* coding sequences, under the control of the CaMV 35S promoter from Cauliflower Mosaic Virus, into the wild-type (Col-0) genetic background, using either the pB7FWG2 vector, in the case of *oeFtsH1-GFP*, *oeFtsH2-GFP* and *oeFtsH8-GFP*, or the pB7RWG2 vector, in the case of *oeFtsH5-RFP*, both purchased by the Flanders Interuniversity Institute for Biotechnology (Gent, Belgium). The *oeFtsH1-GFP gun1-102 ftsh1-1, oeFtsH1*-*GFP gun1-102 ftsh5-3*, *oeFtsH5-RFP gun1-102 ftsh5-3*, *oeFtsH2-GFP gun1-9 ftsh2-3* and *oeFtsH8 gun1-102 ftsh8-1* lines were generated by crossing and PCR-based segregation analysis. Wild-type and mutant seeds were grown on soil in climate chambers under long-day (16 h light/8 h dark) conditions as described by Pesaresi et al. (2009). For lincomycin (Lin) treatments, seeds were surface-sterilized and grown for 6 days (80 μmol m^-1^s^-1^ on a 16h/8h dark/light cycle) on Murashige & Skoog (MS) medium (Duchefa) supplemented with 2% (w/v) sucrose and 1% Phyto-Agar (Duchefa); lincomycin was added at a final concentration of 550 µM, unless otherwise indicated.

### Chlorophyll Fluorescence Measurements and chlorophyll quantification

*In-vivo* Chl a fluorescence was recorded at 6 DAS with a Dual-PAM-100 (Walz) after 20 min of dark adaptation, as described by Pesaresi et al. (2009). Furthermore, the imaging Chl fluorometer (Imaging PAM; Walz) was used to measure i*n vivo* Chl a fluorescence of lincomycin-treated seedlings, according to Tadini et al. (2012). For Chl quantification, 80 mg (fresh weight) of 6-day-old seedlings were ground in liquid nitrogen and extracted in 90% acetone. Chl a and b concentrations were measured according to Porra et al. (1989). Measurements were performed in triplicate.

### Transmission Electron Microscopy

Tissue fragments (1-2 mm^2^) from fully expanded seedlings of Col-0, *gun1-102*, *ftsh2-2*, *ftsh5-3, gun1-102 ftsh5-3* and *gun1-9 ftsh2-3* were fixed in 1,2% glutaraldehyde and 3,3% paraformaldehyde in 0.1 M phosphate buffer (pH 7.4) at 4°C for 2 h, post-fixed in 1% OsO_4_ in the same buffer for 2 h, dehydrated in an ethanol series and embedded in Spurr’s resin. Thin sections (1-2 m thick) were stained with 0.1 % toluidine blue and examined with an Olympus BX-50 light microscope (Olympus, Japan). Ultrathin sections were stained with 2% uranyl acetate and lead citrate, before being examined with a Jeol 100SX TEM (Jeol, Japan) operating at 80 KV.

### Nucleic Acid Analyses

For RNA-blot analyses, total RNA was isolated from 6-day-old Col-0 and mutant seedlings. RNA gel blot analyses were performed according to Meurer et al. (2002), using 5 μg of total RNA for each sample. ^32^P-labeled DNA probes were generated using primers listed in Table S8. For Quantitative Real-Time PCR (qRT-PCR) analyses, 4-µg samples of total RNA, treated with TURBO DNA-free (Ambion by Life Technologies), were employed for first-strand cDNA synthesis, using the GoScript Reverse Transcription system (Promega). qRT-PCR analyses were performed on a CFX96 Real-Time system (Bio-Rad), using primer pairs listed in Table S8. *ACTIN8* (AT1G49240) and *PP2A* (AT1G13320) transcripts were used as internal references. Data obtained from three biological and three technical replicates, were analyzed with the Bio-Rad CFX Manager software (V3.1).

### Expression of recombinant GUN1 and RH50 proteins

The nucleic acid sequences encoding the mature GUN1 and RH50 proteins were cloned into the BamHI-SalI sites of the pMAL-c5X Vector (New England Biolabs). In both cases, the coding sequence of Strep-Tag (WSHPQFEK) was added to the reverse primer (Table S8). Expression of recombinant proteins was performed at 25°C for 4 h with 0,3 mM Isopropil-β-D-1-tiogalattopiranoside (IPTG) and affinity purification was obtained by using the amylose resin affinity matrix (New England Biolabs) in a buffer containing 60 mM HEPES-KOH pH 8.0, 200 mM NaCl, 60 mM KOAc, 10 mM MgOAc supplemented with proteinase inhibitor cocktail (cOmplete™, COEDTAF-RO, Roche).

### Run on transcription assay

Run on transcription assay (Deng et al., 1987) was performed on chloroplasts isolated from 300 mg of 6 DAS Col-0 and *gun1-102* seedlings, grown in presence of lincomycin (Kunst et al., 1998). The transcription reaction was carried out for 10 minutes at 25°C in 50 µl transcription buffer [60 mM HEPES-KOH pH 8.0, 60 mM KOAc, 10 mM MgCl_2_, 10 mM dithiothreitol, 20 RNase inhibitor (Thermo Fisher Scientific), proteinase inhibitor cocktail (cOmplete™, COEDTAF-RO), 150 µM ATP, CTP, GTP, 15 µM UTP and 5 µl [α^32^P]-UTP (3000 µCi/mmol)]. The *rpoA*, *rps12, rbcL^a^*, *rbcL^b^, psaA* and *psbA* DNA probes (2 µg) were blotted onto a Hybond-N + membrane (Amersham) and hybridized with total RNA extracted from chloroplasts. For the *in vitro* complementation of *gun1-102* chloroplasts, 50 ng of GUN1 and RH50 purified recombinat proteins were added to the preparation. Signals were quantified with the ImageJ software.

### Protein sample preparation and immunoblot analyses

For immunoblot analyses, total proteins were extracted from 6-day-old seedlings as described by Martinez-Garcia et al. (1999). Seedling material was homogenized in Laemmli sample buffer (20% [v/v] glycerol, 4% [w/v] SDS, 160 mM Tris-HCl pH 6.8, 10% [v/v] 2-mercaptoethanol) to a concentration of 0.1 mg µl^-1^ (fresh weight/Laemmli sample buffer). Samples were incubated at 65°C for 15 min and, after a centrifugation step (10 min at 16,000 *g*), the supernatant was incubated for 5 min at 95°C and loaded onto SDS-PA gels. For immunoblots to be probed for TIC and TOC proteins (Toc159, Toc75, Toc34, Tic110, Tic40, Tic100, Tic56, Tic20-1), seedling material was homogenized in a modified Laemmli sample buffer (4% [w/v] SDS, 160 mM Tris-HCl pH 6.8, 100 mM dithiothreitol; 0.1 mg µl^-1^ [fresh weight/Laemmli sample buffer]). Samples were incubated at 65°C for 15 min and clarified by a centrifugation step (10 min at 16,000 *g*). The supernatant was then loaded, without denaturation at 95°C. Protein samples corresponding to 4 mg (fw) of seedlings were fractionated by SDS–PAGE (10% [w/v] acrylamide; Schägger and von Jagow, 1987) and then transferred to polyvinylidene difluoride (PVDF) membranes. Replicate filters were immunodecorated with specific antibodies.

Isolation of intact chloroplasts was performed according to Kunst et al. (1998) with minor changes. Aliquots (0.5 g) of 6-day-old seedlings were homogenized in 2 ml of 45 mM sorbitol, 20 mM Tricine-KOH pH 8.4, 10 mM EDTA, 10 mM NaHCO_3_ and 0.1% (w/v) BSA fraction V and centrifuged for 7 min at 700 *g*. The supernatant was collected as the extra-chloroplast fraction, while the pellet was washed with 1 ml of the same buffer. After a centrifugation step (7 min at 700 *g*), the pellet was collected as the intact chloroplast fraction.

Antibodies directed against AtHsp90-1 (AS08 346), Tic40 (AS10 709), Toc34 (AS07 238), Toc75 (AS06 150), ClpB3 (AS09 459), UBQ11 (AS08 307A), ptCpn60 (AS12 2613), cpHsc70-1 (AS08 348), Lhca1 (AS01 005), Lhca2 (AS01 006), Lhca3 (AS01 007), Lhca4 (AS01 008), Lhcb1 (AS01 004), Lhcb2 (AS01 003), Lhcb4 (AS04 045), Lhcb5 (AS01 009) and RbcS (AS07 259) were obtained from Agrisera. The anti-GFP antibody was purchased from Thermo Fisher Scientific, AtHsc70-4 antibody from Antibodies-online, RpoTp (PHY0835S) from PhytoAB, and the anti-RFP antibody was obtained from Chromotek. For Strep-Tag detection, the StrepTactin-HRP conjugated antibody (#1610381) was purchased from Biorad. Tic 100, Tic 56, Tic20-1 antibodies were donated by Masato Nakai (Osaka University), Tic110, Toc159 and ClpC2 antibodies were obtained from Paul Jarvis (University of Oxford), FtsH1 antibody from Zach Adam (The Hebrew University of Jerusalem), and FtsH2 and FtsH5 were donated by Wataru Sakamoto (Okayama University).

### Immunoprecipitation of GUN1-GFP and Toc34

GUN1-GFP immunoprecipitation was performed by using 0,5 g (fresh weight) of 6 DAS seedlings carrying a plastid located GFP protein (cpGFP), used as negative control, *35S::GUN1-GFP* in Col-0 and *35S::GUN1-GFP* in *gun1-102 prpl11-1*. Cotyledons were homogenized in 1 ml of immunoprecitation (IP) buffer (30 mM HEPES-KOH pH 8.0, 200 mM NaCl, 60 mM KOAc, 10 mM MgOAc, 0,5% [v/v] Nonidet P-40 and proteinase inhibitor cocktail [cOmplete™, COEDTAF-RO, Roche]). After a solubilization step (15 min, on ice), samples were subjected to a centrifugation (10 min at 16,000 *g*). DNase treatment was carried out for 1 h at 4°C, by adding 50 U of DNase I (Roche). Supernatants were incubated (2 h, at 4°C) with 20 µl GFP-Trap®_MA (ChromoTek). Beads were then washed 3 times for 10 min with 1 ml of IP buffer and eluted with Laemmli sample buffer. The Toc34 immunoprecipitation was also performed on 0,5 g of frozen Col-0 and *gun1-102* 6 DAS seedlings grown in the presence of 550 µM lincomycin. The powdered tissue was homogenized in 500 µl of IP buffer (30 mM HEPES-KOH pH 8.0, 150 mM NaCl, 60 mM KOAc, 10 mM MgOAc, 1% [v/v] Nonidet P-40 and proteinase inhibitor cocktail [Sigma-Aldrich, P9599]). Samples were incubated on ice for 15 min and subjected to a centrifugation step (10 min at 16,000 *g*). The supernatant was incubated (1 h, at 4°C) with 3 µl of a Toc34-specific antibody. A mixture of magnetic anti-rabbit IgA and IgG Dynabeads™ (Thermo Fisher Scientific) was then added to the samples. After 1 h incubation at 4°C, the magnetic beads were washed 3 times for 10 min with 1 ml of IP buffer and eluted with Laemmli sample buffer.

### BiFC analyses

Vectors were generated by introducing *RPOTp* and *GUN1* coding sequences into pVyNE or pVyCE (Gehl et al., 2009), which encode the N-terminal or the C-terminal portion of the Venus protein, respectively. Transformation of *Agrobacterium tumefaciens*, infiltration of *Nicotiana benthamiana* leaves, and BiFC analyses were performed according to Richter et al. (2013).

### Transient expression and Co-immunoprecipitation of GUN1-GFP and RpoTp-RFP in tobacco leaves

Constructs were generated by introducing *RPOTp* and *GUN1* coding sequences into pB7RWG2 and pB7FWG2 vectors, respectively. Transformation of *Agrobacterium tumefaciens* and infiltration of *Nicotiana benthamiana* leaves were performed as described in Richter et al. (2013). Co-immunoprecipitation assays from total leaf protein extracts and GFP and RFP detection were carried out as described above.

### Soluble total protein extraction

The soluble proteins were extracted from 6-day-old seedlings according as described previously (Kangasjärvi et al., 2008). Seedlings were ground in liquid nitrogen and homogenized in ice-cold extraction buffer (25 mM HEPES-KOH pH 7.5, 10 mM MgCl_2_, protease inhibitor cocktail [complete Mini, Roche], phosphatase inhibitor cocktail [phosSTOP, Roche]). Samples were incubated on ice for 10 min and then centrifuged for 15 min at 16,000 *g*. The supernatant was collected and used for mass spectrometry.

### Mass spectrometry

Identification by data-dependent acquisition (DDA) of the FtsH bands was carried out essentially as described by Konert et al. (2015) and Trotta et al. (2016). The set of doubly and triply charged masses used as the inclusion list for the data-dependent acquisition (DDA) analysis of the bands is reported in Supplemental Table 3. For the analysis of the soluble fraction obtained from total cotyledon extracts, the procedure described for total thylakoid analysis in Trotta et al. (2016) was used, except that 50 μg of protein was used per genotype and treatment, and three biological replicates were digested with trypsin added at a ratio of 1 µg enzyme per 10 µg of sample. The following modifications were also employed: for the nLC-ESI-MS/MS analysis, a single 40 cm x 75 µm column packed with 1.9 µm C18 120 Å (Dr Maisch), and a column oven at 60°C was used (Geyer et al., 2016). The flow rate at 800 bar was 600 nl/min and solvent B consisted of 80% ACN in 0,1% formic acid. The gradient applied was 3 to 43% B for 60 min, followed by an increase to 100% over 5 min, and 10 min of 100% B. The mass spectrometer used was a Q-Exactive HF (Thermo Fisher Scientific), with resolution set to 120000 and scan range 300 to 1800 m/z for MS1 and resolution 15000 with scan range 200 to 2000 m/z for MS2. The dynamic exclusion window was set to 20 s and up to 30 masses with m/z >2+ were selected for MS2 for every MS1 scan. All the other conditions, including the Mascot searches, were conducted as previously described (Trotta et al., 2016), except that all searches have been also repeated allowing for semi-tryptic peptides.

The mass spectrometry analyses performed to study the accumulation of cytosolic chaperones (Hsp90, Hsp70) in the different genetic backgrounds were carried out as follows: differentially 1D-gel lines from Col-0 and gun1-102 were cut, washed in H2O HPLC-grade and dehydrated in 100% ACN for 5 min. After reduction with 25 mM Dithiothreitol (20 min at 56°C) and alkylation with 55 mM of Iodacetamide (20 min at room temperature in the dark) the proteins in the gel slices were digested with trypsin as described by Marsoni et al. (Marsoni et al., 2008). Tryptic peptides were analyzed by liquid chromatography-mass spectrometry (LC-ESI MS/MS). LC-ESI-MS/MS was performed on a Finnigan LXQ linear ion equipped with a Finnigan Surveyor MS pump HPLC system (Thermo Electron Corporation, California, USA). Chromatographic separations were conducted on a Discovery Bio wide-pore C18 column (150 μm I.D. × 150 mm length and 5 μm particle size; SIGMA, USA), using a linear gradient from 5% to 75% CAN in 0.1% formic acid at a flow rate of 2 μL min−1. Acquisitions were performed in the data-dependent MS/MS scanning mode (full MS scan range of 400–1400m/z followed by zoom scan for the most intense ion from the MS scan and full MS/MS for the most intense ion from the zoom scan), thus enabling a dynamic exclusion window of 3 min. Fragmentation spectra were analyzed by MASCOT database search tool (http://www.matrixscience.com/cgi/search_form.pl?FORMVER=2&SEARCH=MISwith) against NCBI protein database (taxonomy restricted to Arabidopsis thaliana) and contaminants database using default settings. Fixed modification of cysteine (carbamidomethylation) and variable modification of methionine (oxidation) was considered. For an accurate protein identification in the Protein family summary significant threshold cut-off was set to 0.01. Only proteins identified with almost two different peptides and protein present in almost three biological replicates were considered for further analysis. Protein relative quantification between Col-0 and gun1-102 sample was done considering the total number of matched peptides assigned to each protein family (the family name was assigned taking into account the top rank hits), normalized using the total number of peptide per gel. Statistical significance was evaluated by T Student test (n=4, p<0.05).

### Statistical analysis

For quantification of immunoblots, RNA blots and plastid precursor protein accumulation, the significance between average values was determined by either the two-tailed paired Student′s t-test, to compare genotypes grown in absence and presence of Lincomycin, and unpaired Student′s t-test, to evaluate different genotypes. Welch’s correction was applied when variances were significantly different, as described by Lemieux et al. (2016). P-values are represented as *, P<0.05; **, P<0.01; ***, P<0.001.

### Accession Numbers

The Arabidopsis Genome Initiative accession numbers for the genes mentioned in this article can be found at TAIR (https://www.arabidopsis.org/): *GUN1* (*AT2G31400*), *FtsH1* (*AT1G50250*), *FtsH2* (*AT2G30950*), *FtsH5* (*AT5G42270*), *FtsH8* (*AT1G06430*), *PRPS21* (*AT3G27160*), *PRPL11* (*AT1G32990*), *cpHsc70-1* (*AT4G24280*), *RH50* (*AT3G06980*), *PP2A* (*AT1G13320*), *RPOTP* (*AT2G24120*), *Tic214* (*ATCG01130*), *rpoA* (*ATCG00740*), *rpoB* (*ATCG00190*), *rps12 3’* (*AtCG00905*/*AtCG01230*), *clpP1* (*AtCG00670*), *atpA* (*ATCG00120*), *psaA* (*ATCG00350*), *rbcL* (*AtCg00490*), *psbA* (*AtCg00020*).

## Acknowledgments

We are grateful to Masato Nakai (Osaka University) for Tic 100, Tic 56, Tic20-1 antibodies, to Paul Jarvis (University of Oxford) for Tic110, Toc159 and ClpC2 antibodies, to Zach Adam (The Hebrew University of Jerusalem) for FtsH1 antibody, and to Wataru Sakamoto (Okayama University) for the FtsH2 and FtsH5 antibodies. *sca3-1* seeds were kindly provided by José Luis Micol. We are also grateful to Msc Azfar Ali Bajwa, Valerio Parravicini, Mario Beretta and Dario Maffi for excellent technical assistance. The Proteomics Facility of the Turku Centre for Biotechnology is acknowledged for their excellent support in mass spectrometry. Academy of Finland grants 307335 and 303757 are acknowledge for financial support. We also thank Paul Hardy for critical reading of the manuscript.

