## Supplemental Figures 1, 2, 3 and Table S1 for "GUN1 influences the accumulation of NEP-dependent transcripts and chloroplast protein import in Arabidopsis cotyledons upon perturbation of chloroplast protein homeostasis"

### Supporting Information

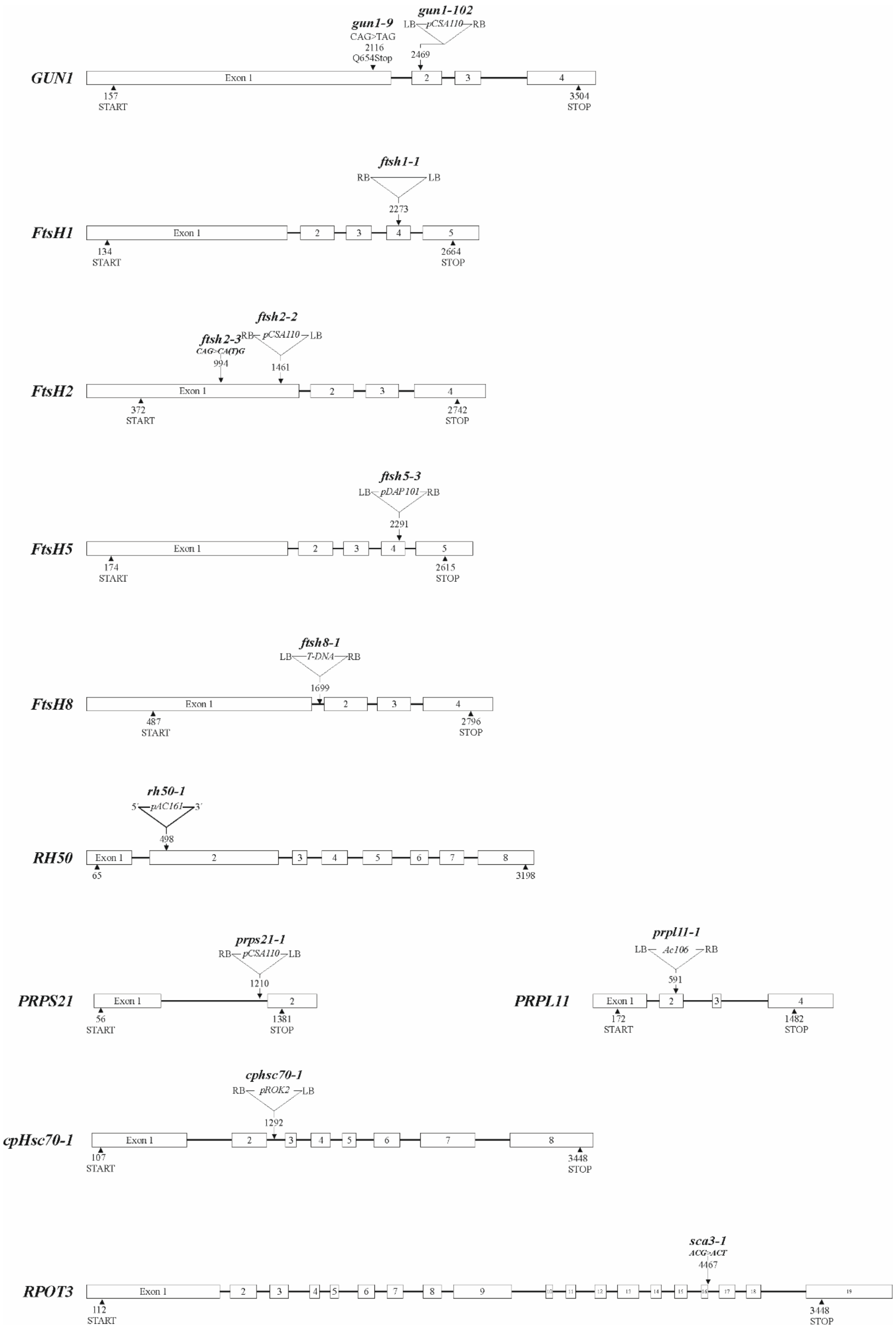

**Figure S1.** Point mutations**,** T-DNA tagging and CRISPR-Cas9 gene editing of *GUN1*, *FtsH*, *RH50*, *PRPS21*, *PRPL11*, *RPOT3* and *cpHSC70-1* genes.

Exons are indicated as numbered white boxes, introns as black lines. Arrowheads indicate the positions of translation initiation and stop codons. Sites, designations and orientations of T-DNA insertions are indicated (RB, right border; LB, left border). The T-DNA insertions are not drawn to scale. Sequence modifications obtained by gene editing are indicated in parentheses: CA(T)G indicates the insertion of a single nucleotide in the first exon of *FtsH2* gene. Note that the phenotype of *gun1-9 ftsh2-3* was fully rescued by the introduction of the wild-type copy of *FtsH2* gene, excluding the possibility of off-target mutations introduced by the CRISPR-Cas9 gene editing strategy.

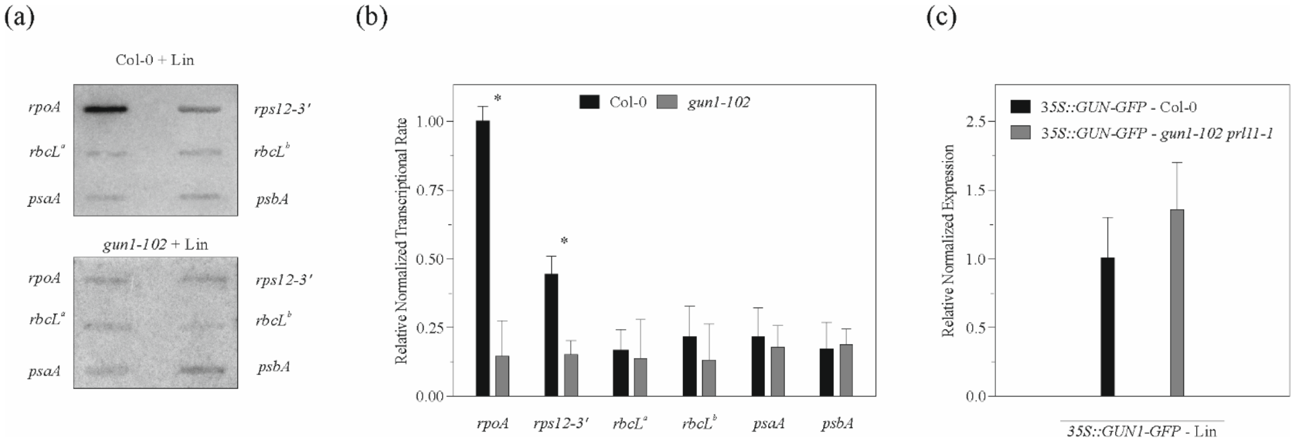


**Figure S2.** Run-on transcription assay in Col-0 and *gun1-102* chloroplasts isolated from seedlings grown on MS medium with Lin and transcript accumulation of the *GUN1-GFP* chimer in Col-0 and *gun1-102 prl11-1* genetic backgrounds. **(a)** Filters with *rpoA*, *rps12-3'* (NEP-dependent genes) and *rbcL*$^a$, *rbcL*$^b$, *psaA*, *psbA* (PEP-dependent genes) probes were hybridized with total [$^{32}$P]-RNA from Col-0 and *gun1-102* isolated plastids. A representative result from three independent experiments is shown. **(b)** Quantification of Run-on signals in (A). Data are the mean ± s.d. obtained from quantification of three independent experiments. Asterisks indicate the statistical significance, as evaluated by Student's t-test ($p<0.05$). **(c)** qRT-PCR analyses to monitor the expression of *35S::GUN1-GFP* construct in Col-0 and *gun1-102 prl11-1* cotyledons grown on MS medium without lincomycin (- Lin). The sequences of the primers used are listed in Table S8. Nine technical replicates (three biological replicates, three technical replicates each) were performed. Lines indicate the Standard deviation (SD). The *ACTIN8* and *PP2A* transcripts were used as internal references.

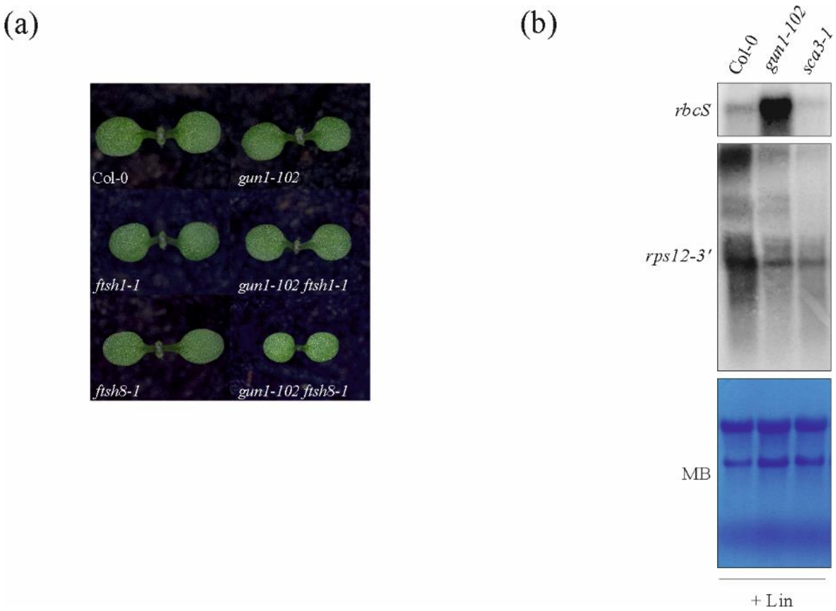


**Figure S3.** Phenotypes of Col-0 and mutant seedlings and norther-blot analyses. **(a)** Phenotypes of Col-0, *gun1-102*, *ftsh1-1*, *ftsh8-1*, *gun1-102 ftsh1-1* and *gun1-102 ftsh8-1* seedlings at 6 DAS, grown on soil under optimal growth-chamber conditions. Note that the same phenotypes were observed when seedlings were grown on MS medium. **(b)** Northern-blot analysis to monitor the expression of the nuclear gene *rbcS* and the plastid NEP-dependent gene *rps12* in Col-0, *gun1-102* and *sca3-1* seedlings. As *gun1-102*, *sca3-1* seedlings are unable to increase NEP-dependent transcript accumulation upon inhibition of PEP activity. Nevertheless, *sca3-1* does not show the *gun* phenotype, as shown by the marked reduction of rbcS transcript accumulation in the presence of Lin.

**Table S1. Quantification of transcripts in light-adapted cotyledons of Col-0 and mutant seedlings grown on MS medium.** Col-0 levels are set to 1 (100%). Values are means ± SD from three independent Northern blots (see Fig. 1). Asterisks indicate the statistical significance, as evaluated by Student's t-test and Welch correction (*, p<0.05; **, p<0.01; ***, p<0.001). [a][Lin] = 11 µM (see Figure 1a); [b][Lin] = 550 µM (see Figure 1a,c,e). The NEP-dependent transcripts are indicated in bold. nd, not detected; na, not analyzed.

| Genotype | *rbcL* | *psaA* | *psbA* | *atpα* | ***Tic214*** | ***rps12-3'*** | ***rpoA*** | ***rpoB*** | ***clpP1*** |
|---|---|---|---|---|---|---|---|---|---|
| *gun1-102* | 1.02 ± 0.06 | 0.95 ± 0.04 | 1.12 ± 0.09 | 0.97 ± 0.06 | 1.09 ± 0.11 | 0.88 ± 0.13 | 0.94 ± 0.08 | 0.93 ± 0.18 | 1.05 ± 0.03 |
| *prps21-1* | 0.75 ± 0.10* | na | 0.54 ± 0.10** | na | na | 3.66 ± 0.50+ | 4.08 ± 0.15*** | na | na |
| *prpl11-1* | 0.52 ± 0.10** | na | 0.31 ± 0.08*** | na | na | 2.58 ± 0.22** | 2.53 ± 0.22*** | na | na |
| *ftsh5-3* | 1.03 ± 0.13 | 0.52 ± 0.06** | 0.41 ± 0.04** | 0.77 ± 0.11 | 1.48 ± 0.10** | 1.67 ± 0.30 | 1.39 ± 0.16* | 1.41 ± 0.28 | 1.38 ± 0.06** |
| *cphsc70-1* | 1.04 ± 0.05 | na | 0.29 ± 0.07*** | na | na | 2.29 ± 0.19** | 1.85 ± 0.26** | na | na |
| *rh50-1* | 0.94 ± 0.13 | na | 1.18 ± 0.08 | na | na | 0.88 ± 0.07+ | 0.94 ± 0.12 | na | na |
| *gun1-102 prps21-1* | 0.81 ± 0.08 | na | 0.31 ± 0.02*** | na | na | 1.39 ± 0.20 | 1.44 ± 0.11** | na | na |
| *gun1-102 prpl11-1* | 0.12 ± 0.02** | na | 0.09 ± 0.03*** | na | na | 0.68 ± 0.07** | 0.29 ± 0.06*** | na | na |
| *gun1-102 ftsh5-3* | 0.78 ± 0.10 | 0.28 ± 0.06*** | 0.34 ± 0.05*** | 0.33 ± 0.09** | 0.98 ± 0.16 | 1.09 ± 0.06 | 0.99 ± 0.16 | 0.98 ± 0.15 | 1.04 ± 0.04 |
| *gun1-102 cphsc70-1* | 0.65 ± 0.04** | na | 0.29 ± 0.09*** | na | na | 0.84 ± 0.13 | 0.89 ± 0.12 | na | na |
| Col-0 + Lin[a] | 0.55 ± 0.10* | na | 0.41 ± 0.12* | na | na | 4.09 ± 0.35** | 2.23 ± 0.13** | na | na |
| Col-0 + Lin[b] | nd | nd | nd | 0.15 ± 0.03** | 8.78 ± 0.17*** | 2.03 ± 0.07** | 4.83 ± 0.37** | 4.27 ± 0.22*** | 3.44 ± 0.04** |
| *gun1-102* + Lin[a] | 0.61 ± 0.06** | na | 0.32 ± 0.04*** | na | na | 1.84 ± 0.05*** | 1.27 ± 0.09* | na | na |
| *gun1-102* + Lin[b] | nd | nd | nd | 0.43 ± 0.11** | 1.89 ± 0.16** | 0.76 ± 0.20 | 0.98 ± 0.10 | 1.28 ± 0.14 | 1.07 ± 0.06 |
| *ftsh5-3* + Lin[b] | nd | nd | nd | 0.06 ± 0.03*** | 3.84 ± 0.25*** | 2.31 ± 0.25* | 4.88 ± 0.11*** | 4.24 ± 0.24*** | 3.79 ± 0.11*** |
| *gun1-102 ftsh5-3* + Lin[b] | nd | nd | nd | 0.13 ± 0.02** | 1.65 ± 0.21* | 0.69 ± 0.10** | 1.03 ± 0.14 | 1.14 ± 0.11 | 0.64 ± 0.11* |
| *rh50-1* + Lin[b] | nd | na | nd | na | na | 2.78 ± 0.17** | 6.94 ± 0.25*** | na | na |

**Table S2.** Quantification of PSMs with medium and high confidences in single replicate injections, identified in total cotyledon soluble extracts from Col-0 and gun1-102 (-/+ Lin). Average and standard deviations, calculated among three biological replicates, are indicated. No statistical differences were revealed among the four groups by the Anova test.

**Table S3.** Identity of proteins present in the four precursor protein bands (hwFtsH) and the four mature protein (mFtsH) bands of Col-0 and *gun1-102* lanes. Total protein extracts from Col-0 and *gun1-102* cotyledons, grown in presence and absence of lincomycin, were fractionated by SDS-PAGE. Two replicate PAGEs were run in parallel; one was blotted, immunodecorated with the FtsH1 antibody, and the film used as a reference to locate and isolate the corresponding precursor regions at 75-90 kDa (hwFtsH band) and the corresponding mature regions at 60-70 kDa (mFtsH band). The Table lists the proteins identified by nLC-ESI-MS/MS on the basis of at least two peptides assigned with at least medium confidence.

**Table S4.** List of masses corresponding to tryptic peptides with charge $2^+$ or $3^+$ predicted from the amino acid sequence of FtsH2 and FtsH5 proteins.

**Table S5.** List of FtsH2 peptides identified by nLC-ESI-MS/MS in the hwFtsH and mFtsH bands of Col-0 (+/- Lin) and *gun1-102* (+/- Lin). Using an inclusion list of targets corresponding to the theoretical masses of the tryptic peptides with charge $2^+$ and $3^+$, generated by the *in silico* digestion of the FtsH2 amino acidic sequence (see Table S4), FtsH2 peptides were identified in all four mFtsH bands and in the hwFtsH band from the Col-0 + Lin sample.

**Table S6**. List of FtsH5 peptides identified by nLC-ESI-MS/MS in the hwFtsH and mFtsH bands of Col-0 (+/- Lin) and *gun1-102* (+/- Lin). Using an inclusion list of targets corresponding to the theoretical masses of the tryptic peptides with charge $2^+$ and $3^+$, generated by the *in silico* digestion of the FtsH5 amino acidic sequence (see Table S3), FtsH5 peptides were identified in all mFtsH and hwFtsH bands.

**Table S7.** Identity of proteins present in the 75 and 90 kDa regions of SDS-PA gels loaded with Col-0 + Lin and *gun1-102* + Lin samples. Gel slices obtained from Col-0 + Lin and *gun1-102* + Lin lanes (see asterisks in Figure 6a) were enzymatically digested with trypsin and analyzed by liquid chromatography-mass spectrometry (LC-ESI MS/MS). Proteins identified in the four gel slices are listed. Data from four biological replicates are reported.

**Table S8.** Sequences of oligonucleotides employed for the molecular characterization of mutant line
